# Circuit-based intervention corrects excessive dentate gyrus output in the Fragile X mouse model

**DOI:** 10.1101/2023.09.27.559792

**Authors:** Pan-Yue Deng, Ajeet Kumar, Valeria Cavalli, Vitaly A. Klyachko

## Abstract

Abnormal cellular and circuit excitability is believed to drive many core phenotypes in fragile X syndrome (FXS). The dentate gyrus is a brain area performing critical computations essential for learning and memory. However, little is known about dentate circuit defects and their mechanisms in FXS. Understanding dentate circuit dysfunction in FXS has been complicated by the presence of two types of excitatory neurons, the granule cells and mossy cells. Here we report that loss of FMRP markedly decreased excitability of dentate mossy cells, a change opposite to all other known excitability defects in excitatory neurons in FXS. This mossy cell hypo-excitability is caused by increased Kv7 function in *Fmr1* KO mice. By reducing the excitatory drive onto local hilar interneurons, hypo-excitability of mossy cells results in increased excitation/inhibition ratio in granule cells and thus paradoxically leads to excessive dentate output. Circuit-wide inhibition of Kv7 channels in *Fmr1* KO mice increases inhibitory drive onto granule cells and normalizes the dentate output in response to physiologically relevant theta-gamma coupling stimulation. Our study suggests that circuit-based interventions may provide a promising strategy in this disorder to bypass irreconcilable excitability defects in different cell types and restore their pathophysiological consequences at the circuit level.

## Introduction

Fragile X syndrome (FXS) is the leading monogenic cause of intellectual disability and autism. This disorder arises from mutations in the *Fmr1* gene resulting in a loss of fragile X messenger ribonucleoprotein (FMRP) (Salcedo-Arellano, Hagerman & Martinez-Cerdeno 2020, Willemsen & Kooy 2017). Hippocampus, the brain area implicated as a central hub for learning and memory, is one of the most affected brain regions in FXS (Bostrom et al 2016). Within the hippocampus, dentate gyrus receives the bulk of cortical inputs and plays a critical role in many core information processing functions (Amaral, Scharfman & Lavenex 2007), which are often dysregulated in neurodevelopmental disorders, including FXS (Deng et al 2022, Eadie et al 2012, Ghilan et al 2015, Lee, Panda & Lee 2020, Yau et al 2019, Yau et al 2016, Yun & Trommer 2011). Distinct from other hippocampal regions, dentate gyrus has two types of glutamatergic excitatory neurons (Figure 1A): granule cells and mossy cells (MCs) (Amaral, Scharfman & Lavenex 2007). Granule cells are the first-station neurons of canonical trisynaptic pathway and the only type of output cells in the dentate gyrus (Amaral, Scharfman & Lavenex 2007); whereas MCs comprise a significant portion of neurons in the hilus region (Scharfman 2016, Scharfman 2018), which is sandwiched between the two blades of the granule cell layer. MCs are in a unique position to control dentate gyrus output because these cells not only can directly excite granule cells, but also indirectly inhibit them through innervating local GABAergic interneurons (Amaral, Scharfman & Lavenex 2007), which play a central role in controlling granule cell activity (Pelkey et al 2017). Both MCs and hilar interneurons have been implicated in various pathological conditions (Pelkey et al 2017, Scharfman 2016, Scharfman 2018, Sloviter et al 2012). However, even though dentate gyrus has attracted extensive attention in the FXS field (Deng et al 2022, Lee, Panda & Lee 2020, Modgil et al 2019, Monday et al 2022, Remmers & Contractor 2018, Sathyanarayana et al 2022, Yau et al 2019), whether or not MC function is abnormal and contribute to hippocampal circuit abnormalities in FXS remains unexplored.

**Figure 1.**
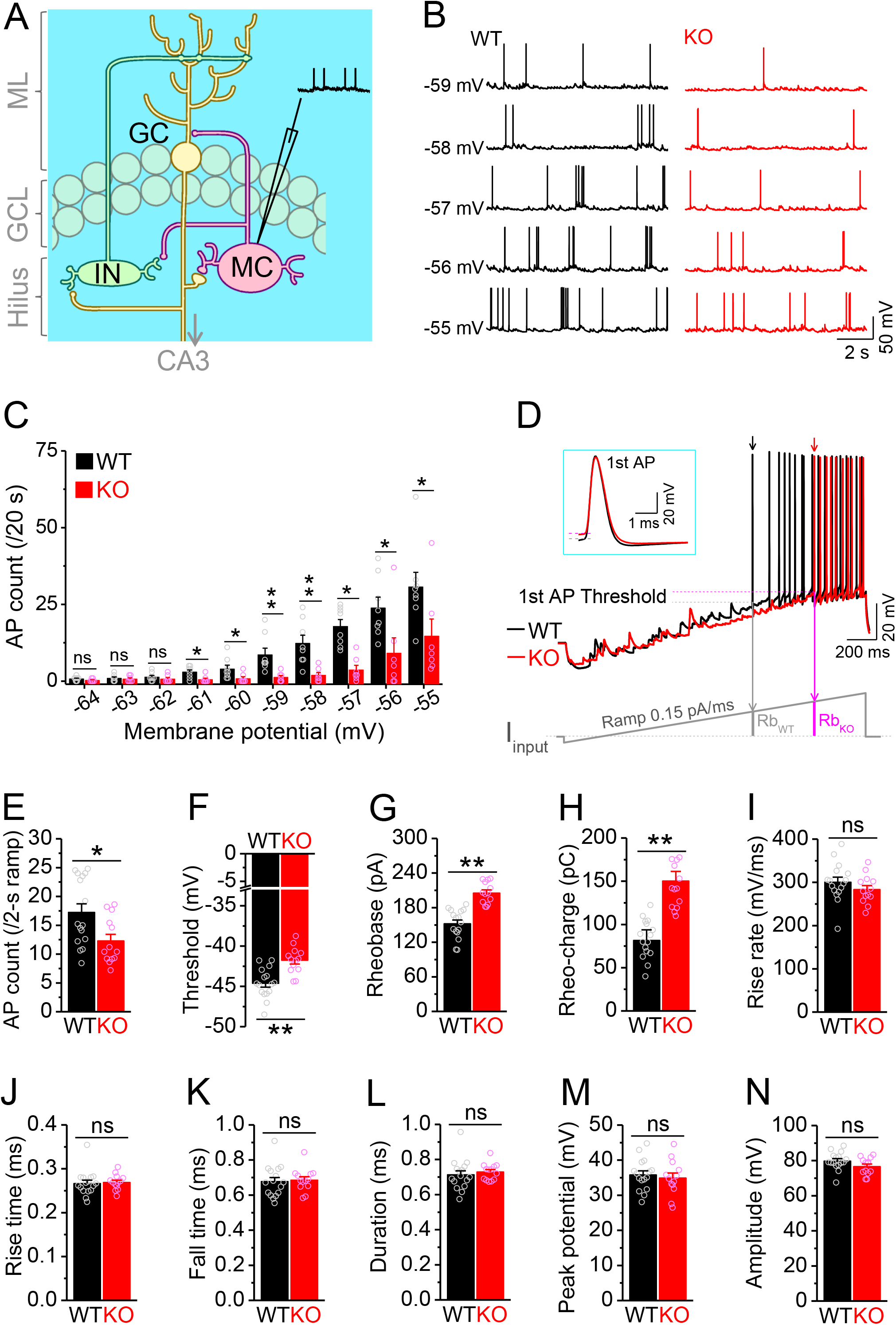
Decreased excitability of dentate mossy cells in *Fmr1* KO mice. (A) Schematic illustration of dentate circuit organization and recordings of APs from mossy cells (MC). Note, for simplicity and clarity, we only show one type of interneurons (IN) with axons terminating onto distal dendrites of a granule cell (GC). The arrow indicates axons of granule cells projecting to CA3. ML, molecular layer; GCL, granule cell layer. (B) Sample traces of spontaneous APs recorded at different membrane potentials from mossy cells in WT (black) and *Fmr1* KO (red) mice. (C) Summary data for experiments exemplified in (A) showing decreased number of APs at membrane potentials of -61 through -55 mV in KO mossy cells. Scatter circles indicate individual data points for this and all subsequent bar graphs in the present study. (D) Determination of AP threshold and rheobase by a ramp current injection (lower trace, ramp rate 0.15 pA/ms). Only the first APs (arrows, which were expanded and aligned by the time of threshold in *insert*) were used to estimate AP parameters. The horizontal lines (*insert*) indicate threshold of the 1st APs. In the *lower* panel, Rb_WT_ and Rb_KO_ denote rheobase current intensity at threshold time-point, and the area (integrating time and input current) enclosed by dotted line, current ramp and Rb_WT_ (or Rb_KO_) are rheobase charge transfer. (D) Phase plots of ramp current-evoked AP traces from (C). Arrows showing the threshold of 1st APs. (E-H) Summary data showing decreased number of APs during 2-s ramp (E), increased voltage threshold (F), rheobase (G) and rheobase charge transfer (H) in KO mossy cells. (I-N) AP upstroke maximum rise rate (I), rise time (J), fall time (K), duration (L), peak potential (M) and amplitude (N). *p < 0.05; **p < 0.01; ns, not significant. The statistical data are listed in Supplementary file 1– Supplementary Table 1. Data are mean ± SEM.

Abnormal neural excitability in FXS has been often linked to dysregulated expression and/or function of various ion channels (Deng & Klyachko 2021), and particularly the family of K^+^ channels, which control neuronal excitability by regulating the action potential (AP) threshold, duration, frequency and firing patterns. Kv7 (KCNQ) channels are a subthreshold-activated and non-inactivating K^+^ conductance that plays key roles in regulating neuronal excitability throughout the brain (Brown & Passmore 2009, Delmas & Brown 2005, Jones, Gamper & Gao 2021, Liu, Bian & Wang 2021), including hippocampal neurons (Gu et al 2005, Incontro et al 2021, Martinello et al 2015, Peters et al 2005). Increasing evidence suggests that Kv7 channels contribute to synaptic plasticity, learning/memory and behavior (Baculis, Zhang & Chung 2020); and their dysfunction contributes to a variety of neurodevelopmental disorders (Baculis, Zhang & Chung 2020, Gilling et al 2013, Jentsch 2000, Jones, Gamper & Gao 2021, Liu, Bian & Wang 2021, Miceli et al 2015, Miceli et al 2008, Nappi et al 2020, Peters et al 2005, Springer, Varghese & Tzingounis 2021). Two members of the Kv7 family (Kv7.2 and Kv7.3) have been identified as targets of FMRP’s translational regulation (Darnell et al 2011), yet it is unknown whether Kv7 expression/function is affected by FMRP loss and plays a role in pathophysiology of FXS.

Here, we report an unexpected hypo-excitability of MCs in FXS mouse model, resulted from upregulation of Kv7 function. We uncover the implications of this defect to dentate dysfunction at the synaptic, cellular, and circuit levels. While this MC excitability defect is unique in being opposite in direction to all other glutamatergic neurons in FXS mouse model, it never-the-less paradoxically leads to an elevated excitation/inhibition (E/I) ratio in granule cells and abnormal circuit processing in the dentate circuit via reduced activity of local inhibitory networks. Our analyses using circuit-wide inhibition of Kv7 channels in *Fmr1* KO mice suggest that circuit-based interventions may represent a promising strategy to bypass the need to correct irreconcilable excitability defects in different cell types, and restore their pathophysiological consequences in this disorder.

## Results

### Decreased excitability of dentate mossy cells in the *Fmr1* KO mice

MCs comprise a large fraction of the cells in the dentate hilar region and are implicated in multiple pathological conditions (Scharfman 2016, Scharfman 2018). To investigate whether MCs are involved in the pathophysiology of FXS, we first examined the excitability of MCs in wildtype (WT) and *Fmr1* knockout (KO) mice, the FXS mouse model. Action potentials (APs) were recorded from MCs at different membrane potentials (from -64 to -55 mV, set by constant current injection). Unexpectedly, we found that loss of FMRP significantly decreased excitability of MCs, as evident by the reduced number of APs fired (Figures 1B-1C; statistical data for every measurement in this study are listed in Supplementary file 1–Supplementary Table 1). This hypo-excitability is opposite to the changes previously observed in other excitatory neurons in FXS models, which typically exhibit hyperexcitability (Contractor, Klyachko & Portera-Cailliau 2015, Deng & Klyachko 2021). To verify this observation, we employed a ramp protocol to evoke APs and examined various AP parameters (Deng et al 2021, Deng et al 2019, Deng et al 2022). Similarly, we found a decreased number of APs fired (Figures 1D-1E), as well as increased voltage threshold, rheobase, and rheobase charge transfer in MCs of *Fmr1* KO mice (Figures 1D and 1F-1H), confirming the hypo-excitable state of MCs in the absence of FMRP. There were no significant changes in AP maximum rise rate, rise time, fall time, duration, peak potential and amplitude in KO MCs (Figures 1I-1N), indicating that the transient Na^+^ current and fast activating K^+^ conductances are likely unaffected in MCs of *Fmr1* KO mice.

The MC hypo-excitability in *Fmr1* KO mice can be attributed to cell-autonomous and/or circuit defects. We first asked whether this defect was caused by changes in the intrinsic passive membrane properties of MCs by examining the resting membrane potential (RMP), cell capacitance and input resistance at RMP level, and found all of them to be unaffected in *Fmr1* KO mice (Figures 2A-2C). However, since input resistance around the threshold level contributes directly to AP initiation, we examined it again at -45 mV (close to threshold level) using a depolarization step in the pharmacologically isolated MCs [using a combination of blockers against both glutamate and GABA receptors (in μM): 10 NMDA, 50 APV, 10 MPEP, 5 gabazine and 2 CGP55845] (Deng et al 2019, Deng et al 2022), as well as in the presence of 1 μM TTX and 10 μM CdCl_2_ to block Na^+^ and Ca^2+^ channels, respectively. Under these conditions, the input resistance was significantly lower in *Fmr1* KO than WT MCs (Figure 2D-2E), suggesting that the reduced input resistance around the threshold potential may be a primary contributor to MC hypo-excitability in *Fmr1* KO mice.

**Figure 2.**
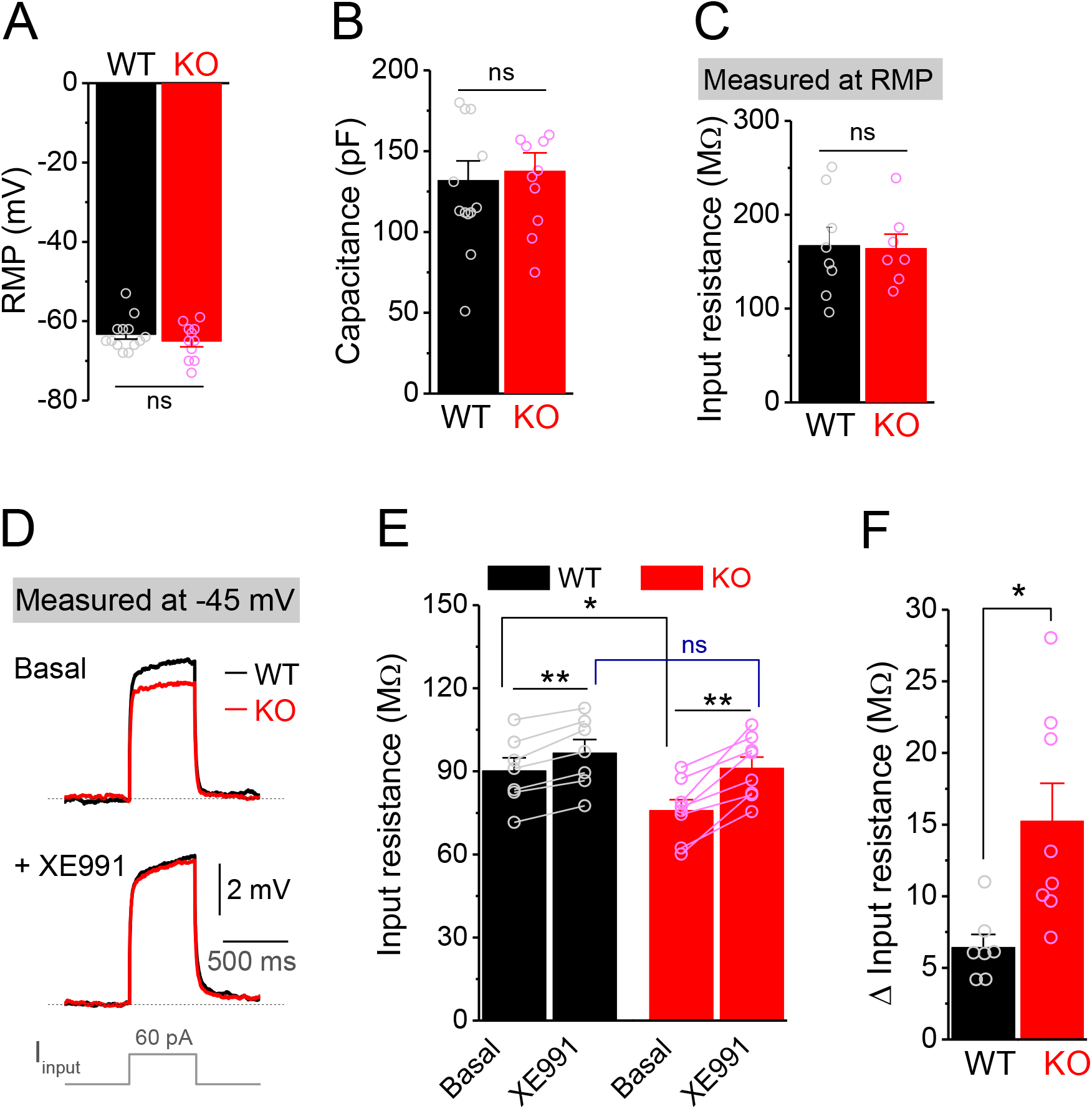
Decreased input resistance around threshold potential in *Fmr1* KO mossy cells. (A and B) Resting membrane potential (RMP, A) and membrane capacitance (B) of mossy cells. (C) Input resistance measured at RMP level. (D) Input resistance measured at -45 mV. Sample traces of the depolarization current step (*lowermost panel*) induced voltage responses before (Basal, *upper panel*) and during XE991 (+ XE991, *middle panel*). (E) Summary data of input resistance before (basal) and during XE991. (F) Effects of XE991 on increasing of input resistance. Note XE991 have stronger effect on KO mossy cells. *p < 0.05; ns, not significant. The statistical data are listed in Supplementary file 1– Supplementary Table 1. Data are mean ± SEM.

### Enhanced Kv7 function causes mossy cell hypo-excitability in *Fmr1* KO mice

Among conductances active at sub-threshold potentials, Kv7 channels are known to play a major role in determining the input resistance (Baculis, Zhang & Chung 2020, Greene & Hoshi 2017, Jones, Gamper & Gao 2021, van der Horst, Greenwood & Jepps 2020). We thus tested whether the reduced input resistance in *Fmr1* KO MCs is associated with Kv7 dysfunction. We found that the Kv7 blocker XE991 (10 μM) increased input resistance in all tested MCs from both WT and *Fmr1* KO (Figure 2E), with a significantly larger effect in KO mice (Figure 2F). Importantly, XE991 abolished the difference in input resistance at -45 mV between WT and *Fmr1* KO cells (Figure 2E). Moreover, Kv7 inhibition with XE991 also caused a larger shift in the holding current in KO than WT MCs (Figure 3A), and a larger membrane depolarization in KO than WT MCs when the membrane potential was initially set at -45 mV (Figure 3B). We further observed that loss of FMRP increased Kv7 currents in MCs as measured by a ramp protocol (Figures 3C-3D). Together, these results strongly suggest that Kv7 function is increased in *Fmr1* KO MCs.

**Figure 3.**
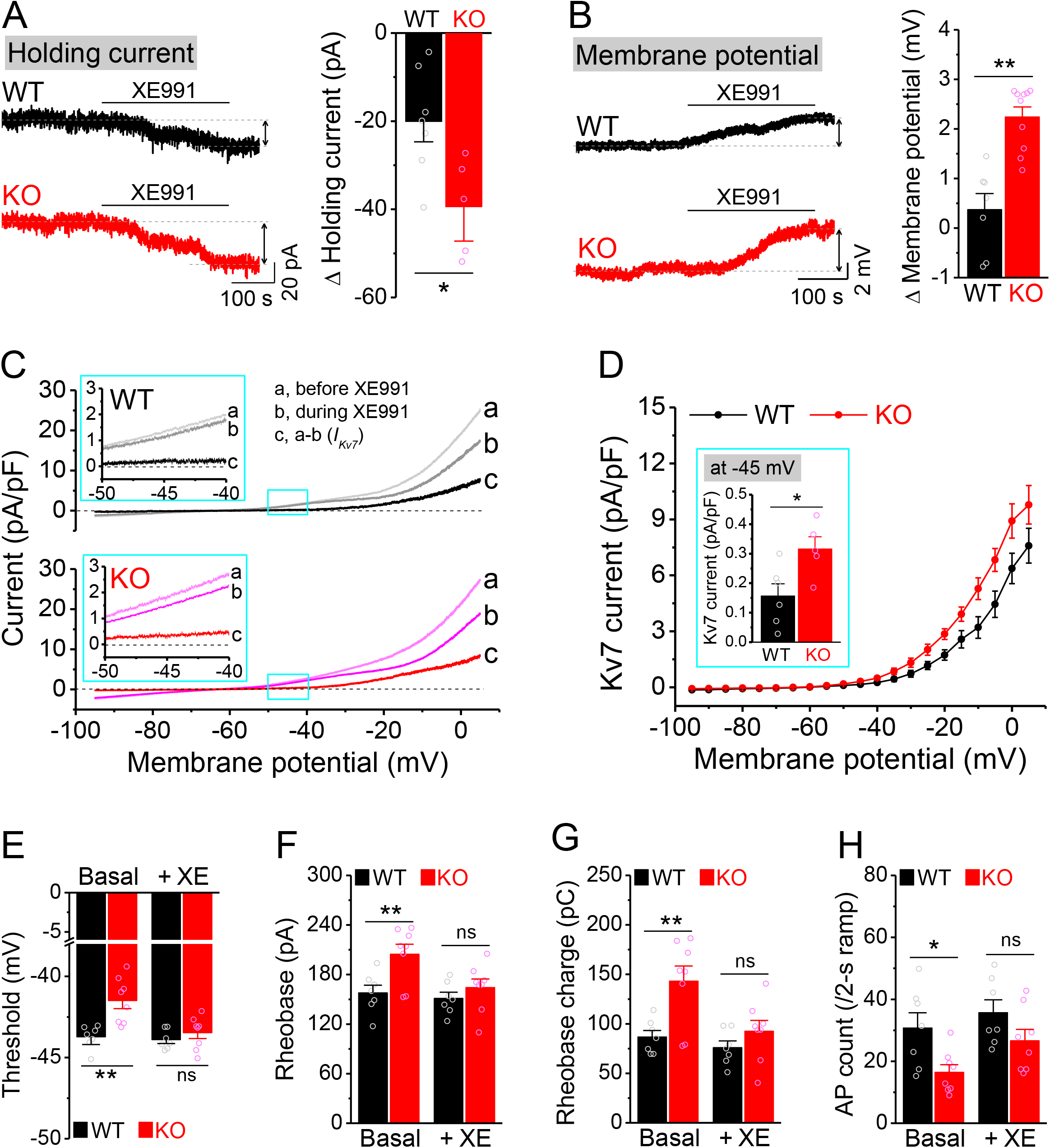
Enhanced Kv7 function causes hypo-excitability of *Fmr1* KO mossy cells. (A) Changes in holding current at -45 mV in response to XE991. *Left*, sample traces; *Right*, summary data. (B) Changes in membrane potential in response to XE991 when the initial potential being set at -45 mV. *Left*, sample traces; *Right*, summary data. (C) Kv7 current was induced by a ramp protocol (from -95 to +5 mV with a rate of 0.02 mV/ms) and determined by XE991 sensitivity. *Inserts*, the enlargements of boxed areas in main traces. (D) The I-V curves were constructed from ramp-evoked Kv7 currents every 5 mV (quasi-steady-state current, averages over 0.01 mV intervals) and normalized to respective cell capacitances. Insert, Kv7 current at -45 mV. (E-H) Increased threshold (E), rheobase (F) and rheobase charge transfer (H), as well as decreased number of APs (H) in pharmacologically isolated KO MCs. XE991 abolished these differences between genotypes. *p < 0.05; **p < 0.01; ns, not significant. The statistical data are listed in Supplementary file 1– Supplementary Table 1. Data are mean ± SEM.

The ‘classical’ Kv7 current is carried predominantly by heteromeric KCNQ2/KCNQ3 channels (Brown & Passmore 2009). KCNQ2 and KCNQ3 have been identified as targets of FMRP translational regulation (Darnell et al 2011) and are highly expressed in the hippocampus (Tzingounis et al 2010). We thus examined if KCNQ2 and KCNQ3 levels were altered by FMRP loss, but found no changes in expression of either isoform in the whole brain lysate as measured by Western blot (Figure 3—figure supplement 1A-1B). To exclude the possibility that expression variation among brain regions masked KCNQ changes in a particular area, we performed the same experiment on dentate gyrus lysate only. We found no detectable changes in expression of KCNQ2 or KCNQ3 in *Fmr1* KO mice (Figure 3—figure supplement 1C-1D). We did not further investigate the precise mechanisms underlying enhancement of Kv7 function in the absence of FMRP, since the present study primarily focuses on the functional consequences of abnormal cellular and circuit excitability.

To verify the role of elevated Kv7 function in decreased excitability of *Fmr1* KO MCs, we first confirmed that hypo-excitability is still observed in these cells when they were isolated from circuit activity using blockers of both glutamate and GABA receptors (as above). We found this to be the case, as evident by increased threshold, rheobase and rheobase charge transfer (Figures 3E-3G), as well as decreased number of APs (Figure 3H) in circuit-isolated MCs of *Fmr1* KO comparing to WT mice. Importantly, Kv7 blocker XE991 abolished all these differences in excitability metrics between genotypes (Figures 3E-3H).

Together, these results indicate that the hypo-excitability of MCs in *Fmr1* KO mice has primarily a cell-autonomous origin and can be attributed to abnormally elevated Kv7 function.

### Mossy cell hypo-excitability dominates adaptive circuit changes in *Fmr1* KO mice

Next we examined if, in addition to cell-autonomous Kv7-mediated defects, there is also a circuit contribution to MC hypo-excitability. First, we measured spontaneous and miniature excitatory postsynaptic currents (sEPSCs and mEPSCs), and found that the excitatory drive onto MCs was increased, as evident by the elevated number and instantaneous frequency of both sEPSCs and mEPSCs in KO MCs (Figures 4A, 4B, Figure 4—figure supplement 1A-1B), without changes in their amplitudes (Figures 4E and Figure 4—figure supplement 1C). We note that this increase in excitatory drive cannot explain the MCs’ hypo-excitability; rather these results likely reflect the hyperexcitable state of granule cells (Deng et al 2022), which provide the majority of excitatory drive onto MCs. We next examined inhibitory inputs onto MCs and found that the number and instantaneous frequency of spontaneous inhibitory postsynaptic currents (sIPSCs) was also increased in KO MCs (Figures 4C-4D) without changes in amplitude (Figure 4F). However, we did not observe significant differences in miniature inhibitory postsynaptic currents (mIPSCs) in KO MCs (Figure 4—figure supplement 1D-1F). Because the overall circuit contribution to MC excitability is determined by the net effect of both excitatory and inhibitory drives, we examined the E/I ratio of these inputs, a parameter which integrates changes in both excitatory and inhibitory drives into a single variable and thus allows us to determine the net circuit effect on MCs excitability. Given that the amplitudes of sEPSCs and sIPSCs were indistinguishable between genotypes (Figures 4E-4F), we could estimate the E/I ratio simply by using the mean frequency of spontaneous synaptic events (Figures 4G-4H), and observed an increased E/I ratio in *Fmr1* KO MCs (↑ ∼20%, Figure 4I) which acts in opposition to the intrinsic hypo-excitability observed in MCs. Thus these adaptive circuit changes likely function as a compensatory mechanism to the intrinsic MCs’ hypo-excitability in *Fmr1* KO mice, similarly to adaptive circuit changes also seen in the cortex (Antoine et al 2019). However, this circuit adaptation is insufficient to compensate for MC excitability defects because we observed overall hypo-excitability in *Fmr1* KO MCs in the intact circuits.

**Figure 4.**
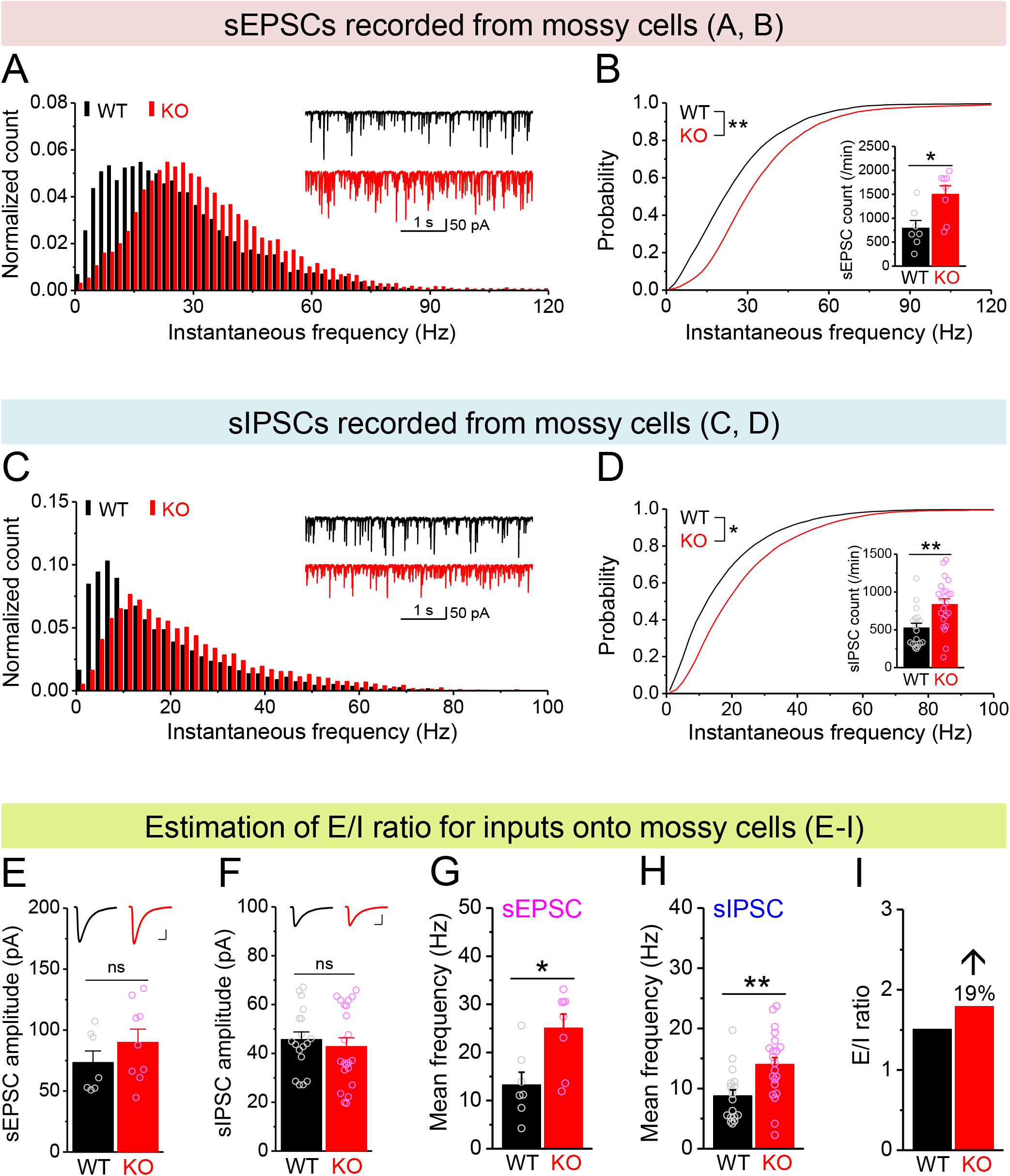
Increased E/I ratio of inputs onto *Fmr1* KO mossy cells. (A) Distribution of sEPSC instantaneous frequency in MCs. A bin size of 2 Hz was used to calculate sEPSC frequency distribution from a 30-s-long trace per cell. The number of sEPSCs within each bin was normalized to the total number of the respective cells for pooling the data from all cells. Note that sEPSC events in KO mice had a shift toward high frequency. *Insert*, sample traces of sEPSCs for WT (black) and KO (red) mice. (B) Cumulative probability of sEPSC instantaneous frequency in MCs. *Bar graph*, number of sEPSCs per minutes. Note that both cumulative probability and number of sEPSCs reveal increased excitatory drive onto MCs. (C and D) sIPSCs recorded from KO and WT MCs, aligned in the same way as in (A and B), respectively. The IPSC signals were down-ward here and also in Figures 5, Figure 4—figure supplement 1 and Figure 5—figure supplement 1, due to a high chloride electrode solution was used in these experiments. Note the increased sIPSC frequency and number in KO MCs. (E and F) Summary data for sEPSC amplitude (E) and sIPSC amplitude (F) recorded from MCs. *Inserts,* sample sEPSC (E) and sIPSC (F) events for WT (black) and KO (red) MCs. Scale: 5 ms (horizontal) and 25 pA (vertical). (G and H) Mean frequency of sEPSCs (G) and sIPSCs (H) recorded from MCs. Note that loss of FMRP increased mean frequency of both sEPSC and sIPSC. (I) E/I ratio evaluated by sEPSC and sIPSC frequencies (mean values from G and H, respectively). Note the increased E/I ratio in *Fmr1* KO mice. *p < 0.05; **p < 0.01; ns, not significant. The statistical data are listed in Supplementary file 1– Supplementary Table 1. Data are mean ± SEM.

### Mossy cell defects reduce excitatory drive onto hilar interneurons in *Fmr1* KO mice

The principal output of the dentate circuit is determined by granule cell firing, which is largely controlled by the balance of the excitatory inputs to granule cells from the stellate cells of the entorhinal cortex via the perforant path (PP), local excitatory inputs from MCs and inhibitory inputs from local interneurons. In the *Fmr1* KO mice, stellate cells have normal excitability (Deng & Klyachko 2016); the granule cells are hyper-excitable (Deng et al 2022); while MCs are hypo-excitable. We thus next probed excitability of the hilar interneurons, the remaining component of this circuit that has not been examined thus far. Because these inhibitory neurons exhibit both morphological and electrophysiological diversity and no clear correlation between morphology and electrophysiological properties has been observed (Mott et al 1997), here we classified a total of 66 recorded interneurons into 3 types according to AP firing pattern: 1) fast-spiking interneurons (high-frequency non-adapting firing); 2) regular-spiking interneurons (slower and adapting firing); and 3) stuttering-like interneurons (high-frequency irregular bursting firing) (Golomb et al 2007) (Figure 5—figure supplement 1A). Chi-square test showed no significant differences in the ratios of the 3 types of interneurons between WT and KO mice (Figure 5—figure supplement 1A). Furthermore, we found that the passive membrane properties (RMP, capacitance and input resistance) and threshold of these interneurons were not significantly different between genotypes and among cell types (thus data were pooled, Figure 5—figure supplement 1B-1E). These results indicate that the intrinsic excitability of hilar interneurons is not affected significantly in the absence of FMRP, suggesting that alterations in synaptic drive onto these cells may play a critical role in determining changes in local inhibition in *Fmr1* KOs.

Therefore, we next examined excitatory and inhibitory drives onto hilar interneurons by recording spontaneous and miniature synaptic inputs to these cells. We found that the number and instantaneous frequency of sEPSCs and mEPSCs were markedly decreased in *Fmr1* KO mice (Figures 5A, 5B, and Figure 5—figure supplement 1F-1G) with no changes in amplitudes (Figures 5E and Figure 5—figure supplement 1H). Further, the number and instantaneous frequency of sIPSCs were also decreased in KO interneurons (Figures 5C-5D) without change in amplitude (Figure 5F), while no significant changes in mIPSCs was observed (Figure 5— figure supplement 1I-1K). We then estimated the balance of E/I inputs onto interneurons using the same approach as above for MCs and observed a markedly decreased E/I ratio of inputs onto *Fmr1* KO interneurons (↓ ∼60%, Figure 5I). We note that changes in the excitatory drive onto interneurons include both mEPSC and sEPSC frequencies, which reflect not only potential deficits in excitability of their input cells, such as MCs, but also changes in synaptic connectivity/function, that may arise from homeostatic circuit reorganization/compensation (see Discussion).

**Figure 5.**
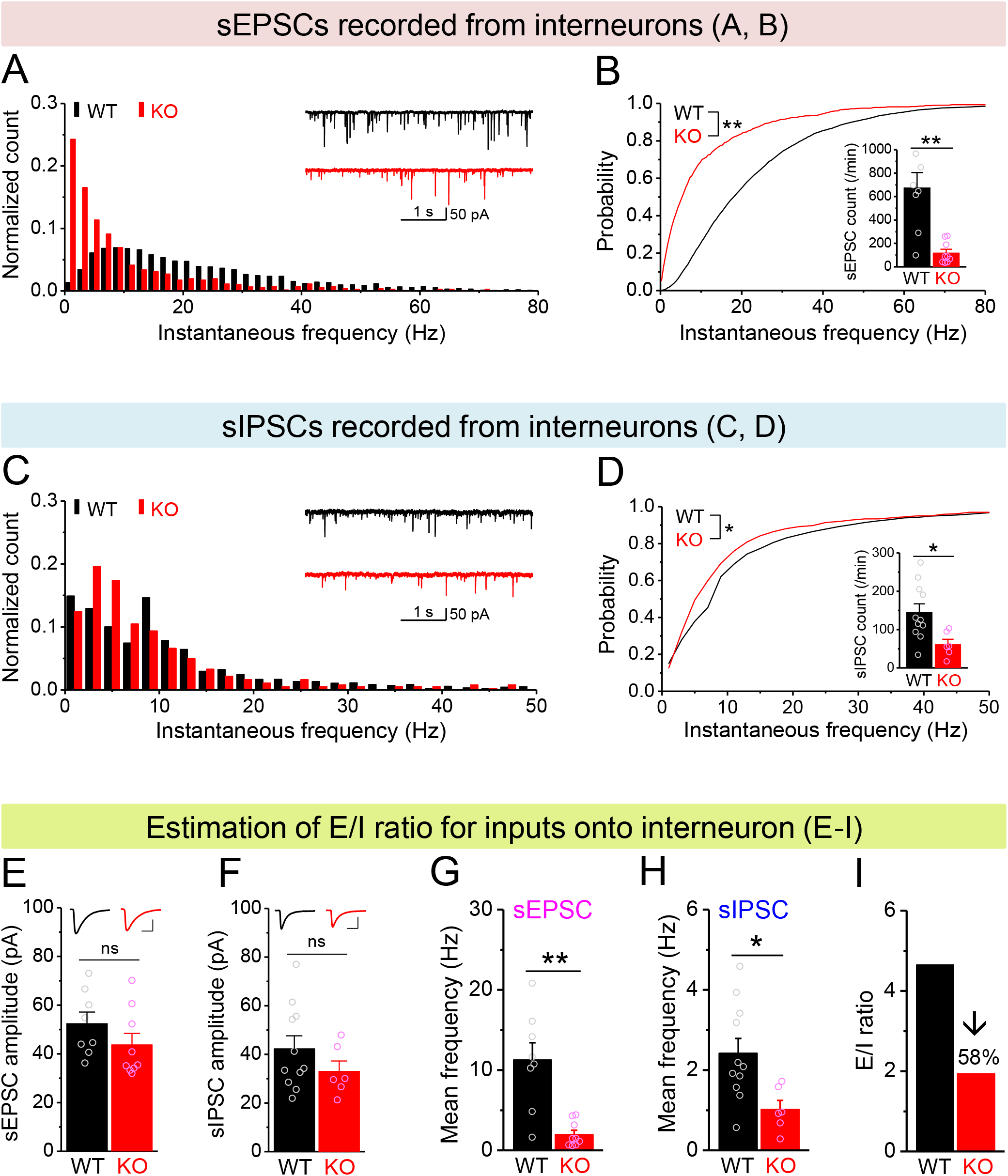
Decreased E/I ratio of inputs onto *Fmr1* KO hilar interneurons. (A) Distribution of sEPSC instantaneous frequency in hilar interneurons. Note that KO sEPSC events had a shift toward low frequency. *Insert*, sample traces of sEPSCs for WT (black) and KO (red) interneurons. (B) Cumulative probability of sEPSC instantaneous frequency in interneurons. *Bar graph* shows number of sEPSCs per minutes. Note that both cumulative probability and number of sEPSCs reveal decreased excitatory drive onto interneurons. (C and D) sIPSCs recorded from interneurons, aligned in the same way as in (A and B), respectively. Note the decreased sIPSC frequency and number in KO interneurons. (E and F) Summary data for sEPSC amplitude (E) and sIPSC amplitude (F) recorded from interneurons. *Inserts,* sample sEPSC (E) and sIPSC (F) events for WT (black) and KO (red) MCs. Scale: 5 ms (horizontal) and 25 pA (vertical). (G and H) Mean frequency of sEPSCs (G) and sIPSCs (H) recorded from interneurons. Note loss of FMRP decreased mean frequency of both sEPSC and sIPSC. I) E/I ratio evaluated by sEPSC and sIPSC frequencies in interneurons (mean values from G and H, respectively). Note the decreased E/I ratio in *Fmr1* KO mice. *p < 0.05; **p < 0.01; ns, not significant. The statistical data are listed in Supplementary file 1– Supplementary Table 1. Data are mean ± SEM.

Considering that excitability of stellate cells is largely unaffected and granule cells show profound hyperexcitability in *Fmr1* KO mice, our observations suggest that it is the MCs that provide the dominant excitatory drive to hilar interneurons, causing a marked decrease in E/I ratio of inputs onto interneurons in *Fmr1* KO mice. To determine if this is indeed the case, we took advantage of different properties of PP, granule cell and MC axonal terminals (Chancey et al 2014, Chiu & Castillo 2008, Shigemoto et al 1997). Specifically, granule cell and PP axonal terminals contain group II mGluRs (Shigemoto et al 1997), while MC axonal terminals express type I cannabinoid (CB1) receptors (Chancey et al 2014, Chiu & Castillo 2008). Accordingly, mGluR Group II agonist DCG-IV (1 μM) selectively inhibits granule cell-and PP-derived EPSCs onto hilar interneurons; while the CB1 agonist WIN 55212-2 (WIN, 5 μM) selectively inhibits MC-derived EPSCs onto these interneurons. In this analysis, for simplicity and better comparison among cells, we normalized the rate of sEPSCs to its own baseline before agonist application (i.e., normalized frequency). We observed that DCG-IV had little effect on the normalized frequency of sEPSCs recorded in interneurons in both genotypes (Figures 6A, 6C, 6D, 6F), indicating that granule cell-and PP-derived EPSCs comprise a limited portion of excitatory drive onto hilar interneurons in both genotypes. As a control for DCG-IV effectiveness, we observed that DCG-IV strongly reduced the normalized frequency of sEPSCs recorded from MCs that primarily receive excitatory inputs from granule cells (Figures 6G-6I). In contrast, even though WIN markedly reduced the normalized frequency of sEPSCs in interneurons of both WT and KO mice (Figures 6B, 6C, 6E and 6F), it failed to affect the normalized frequency of sEPSC recorded from mossy cells (Figures 6H, 6I), suggesting that MCs provide a significant proportion of excitatory drive onto hilar interneurons. Together with the observations above, this analysis supports the notion that the hypo-excitability of MCs in *Fmr1* KO mice is a major factor contributing to the reduction of excitatory drive onto hilar interneurons, which ultimately results in reduced local inhibition.

**Figure 6.**
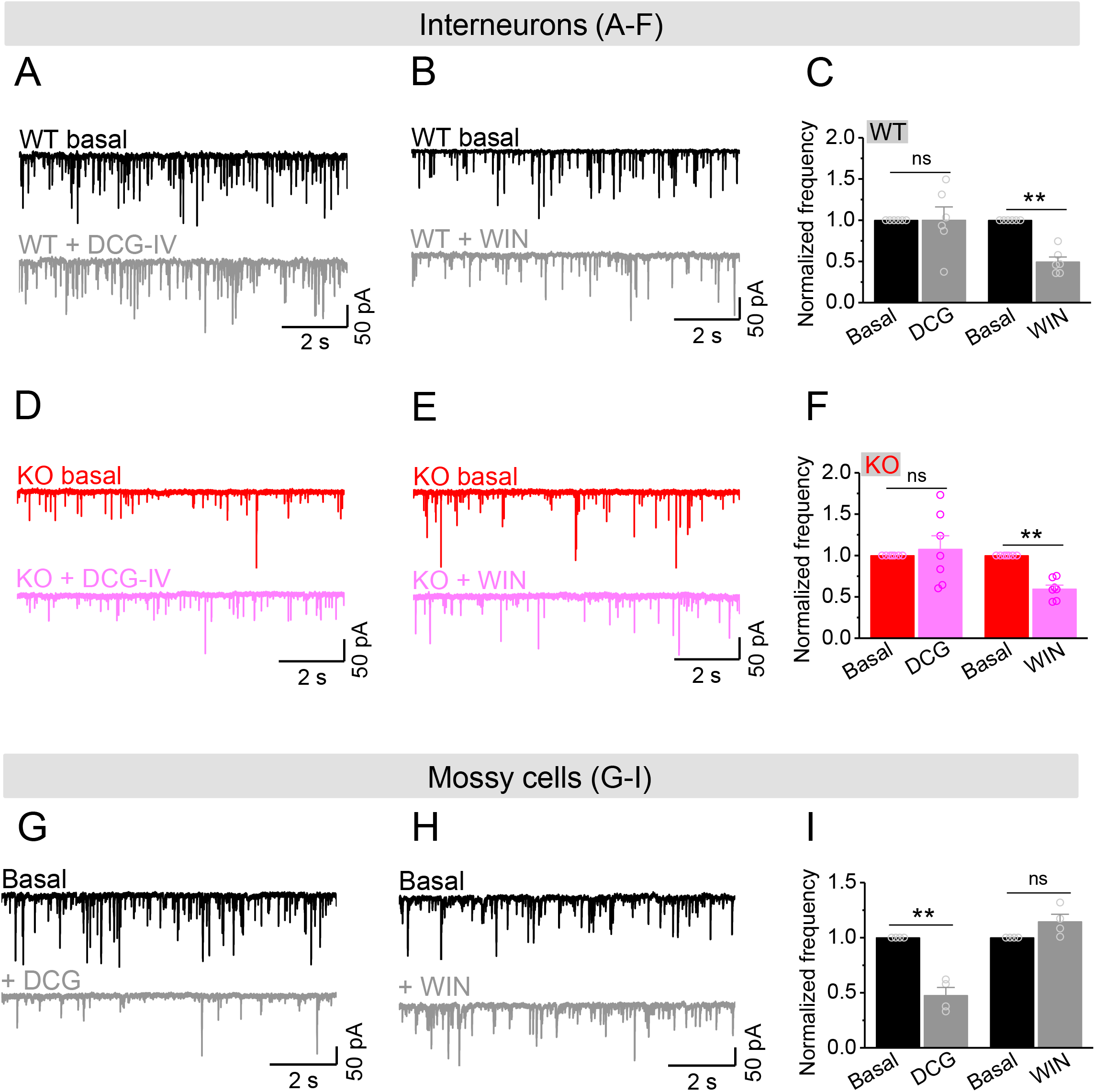
Mossy cells provide the main excitatory drive onto hilar interneurons. (A) Sample traces of sEPSCs recorded from an interneuron of WT mouse before and during DCG-IV. (B) The same as in (A), but for WIN55212-2 in an interneuron of WT mouse. (C) Effect of DCG-IV or WIN55212-2 on the sEPSC normalized frequency recorded from interneurons of WT mice. (D-F) The same as in (A-C) but for interneurons of KO mice. Note that WIN55212-2 had comparable effects on normalized frequency of sEPSCs in KO and WT interneurons, but DCG-IV did not have measurable effects on both genotypes. (G) Control experiment showing effectiveness of DCG-IV. Sample traces of sEPSCs recorded from MCs before and during DCG-IV. (H) Sample traces of sEPSCs recorded from MCs before and during WIN 55212-2. (I) Summary data of changes in the normalized frequency of MC sEPSCs in response to DCG-IV (47.6 ± 7.2% of basal) and WIN55212-2 (114.5 ± 6.7% of basal). Note that, compared to interneurons (A-F), MCs exhibited opposite response to these two agonists, indicating the effectiveness of both agonists. **p < 0.01; ns, not significant. The statistical data are listed in Supplementary file 1– Supplementary Table 1. Data are mean ± SEM.

### Circuit-wide inhibition of Kv7 channels increased local inhibitory drive and corrected granule cell excitability in *Fmr1* KO mice

If upregulation of Kv7 function is largely limited to MCs in the dentate circuit of *Fmr1* KO mice, then our findings suggest that inhibition of Kv7 channels would be more efficient to enhance MC excitability in KO than WT mice and thus may be more effective to boost the inhibitory drive onto *Fmr1* KO granule cells, thus reducing granule cell hyperexcitability in KO mice. To test this hypothesis, we first simultaneously recorded sEPSCs and sIPSCs in the granule cells to get a better quantitative analysis of the E/I input onto these cells. Granule cells were held at -40 mV resulting in sEPSC appearing as a down-ward current, while sIPSC appearing as up-ward current, which was verified by selective blockers of GABA_A_ or AMPA and NMDA receptors (Figure 7—figure supplement 1A). We found that loss of FMRP did not affect the mean frequencies, amplitudes and charge transfer of sEPSCs and sIPSCs onto granule cells (Figures 7B, 7C and Figure 7—figure supplement 1B-1G), and thus did not alter the baseline E/I ratio as determined by any of these parameters (Figures 7D, Figure 7—figure supplement 1D-1G). In line with our hypothesis, circuit-wide inhibition of Kv7 channels with bath application of XE991 markedly increased sIPSC frequency in the granule cells of *Fmr1* KO but not in WT mice (Figure 7C), while having no effect on sEPSC frequency in either genotype (Figure 7B). As a result, XE991 application shifted the E/I balance toward stronger inhibition in the *Fmr1* KO, but not in WT mice (Figure 7D). These results indicate that circuit-wide inhibition of Kv7 channels could be effective in reducing granule cell hyperexcitability in the KO mice. We noted that XE991 also slightly reduced both sEPSC and sIPSC amplitudes, but to a similar extent in both genotypes (Figure 7—figure supplement 1B-1C), and without detectable changes in charge transfer (Figure 7—figure supplement 1E-1F).

**Figure 7.**
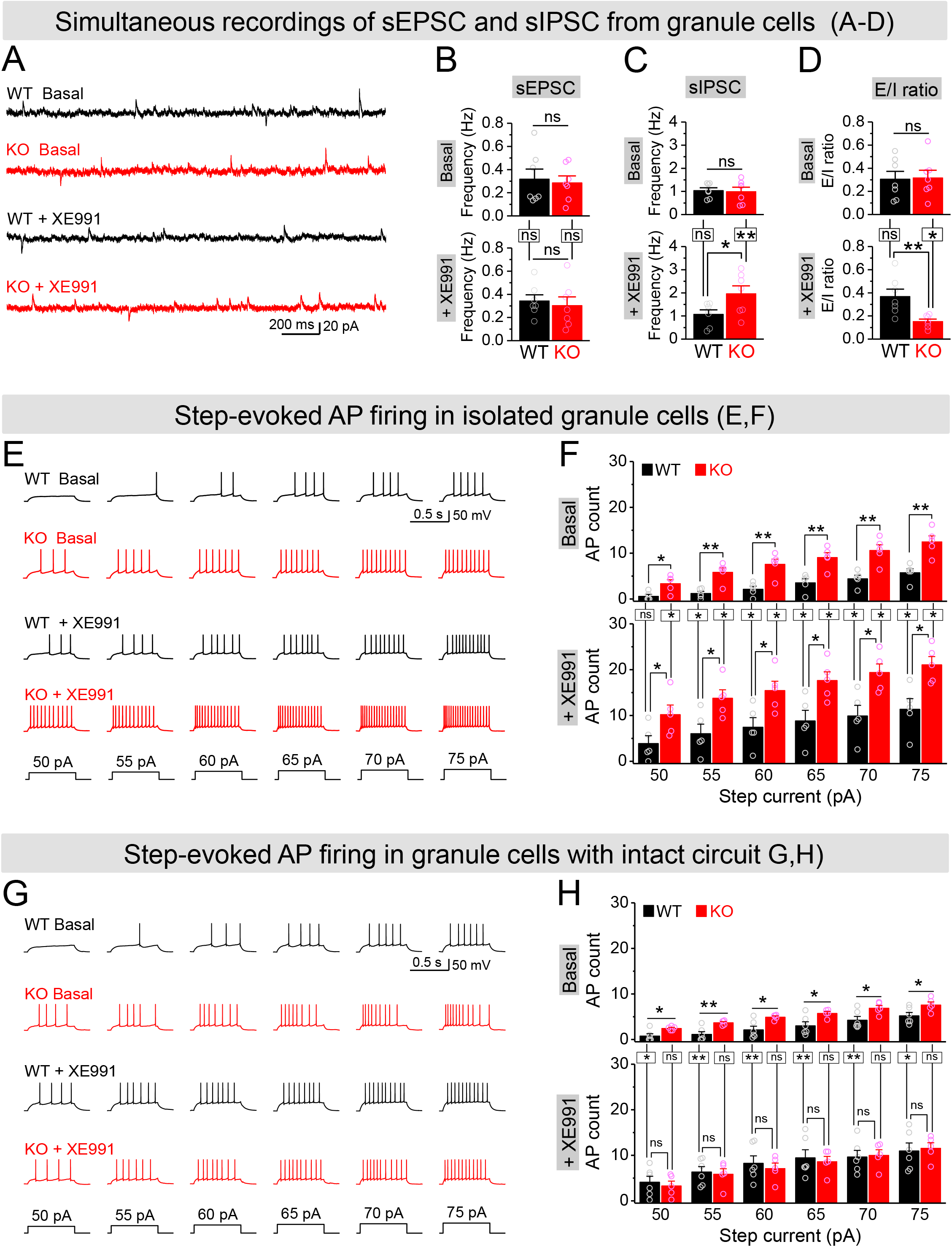
Circuit-wide inhibition of Kv7 channels boosted inhibitory drive onto granule cells and corrected granule cell excitability in *Fmr1* KO mice. (A) Sample traces of simultaneous recording of sEPSC (down-ward events) and sIPSC (up-ward events) from granule cells (also see Figure 7—figure supplement 2A). (B) Summary data for sEPSC mean frequency in basal (*upper*) and during XE991 (*lower*). Horizontal lines (with or without dropdown) denote comparison between genotypes; vertical lines indicate comparison between before and during XE991 within genotypes (C) The same as in (B), but for sIPSC simultaneously recorded from the same granule cells. Note that XE991 increase sIPSC frequency only in KO mice, but not in WT mice. (D) E/I ratio evaluated by frequency. XE991 (*lower*) significantly decreased the E/I ratio in KO mice only. (E) Evaluation of granule cell excitability by recording APs. Sample traces for multistep-current (*Lowermost panel*) evoked APs in WT and KO granule cells in the pharmacologically isolated granule cells in the absence (*upper*) or presence of XE991 (*lower*). (F) Summary data for number of APs exemplified in (E) showing increased excitability of the pharmacologically isolated granule cells in the absence (*upper*) or presence of XE991 (*lower*). Also, note that XE991 increased number of AP in both WT and KO granule cells. Horizontal lines (with dropdown) denote comparison between genotypes; vertical lines indicate comparison between in the absence and presence of XE991 within genotypes. (G) The same as in (E), but for granule cells with intact dentate circuit. (H) The same as in (F), but for granule cells with intact circuit. Note that XE991 did not increase number of AP in KO granule cells; rather it increased number of AP only in WT granule cells. *p < 0.05; **p < 0.01; ns, not significant. The statistical data are listed in Supplementary file 1– Supplementary Table 1. Data are mean ± SEM.

To examine more directly if circuit-wide inhibition of Kv7 channels could be effective in reducing granule cell hyperexcitability in the KO mice, we performed analyses of granule cell firing – the ultimate measure of cellular excitability. APs were evoked by a multistep current injection (55–75 pA, Figure 7E). To disentangle the direct-and circuit-mediated-actions of Kv7 inhibition, we first synaptically isolated granule cells from the circuit using the same pharmacological approach as described above, and observed that loss of FMRP strongly increased the AP firing in granule cells (Figures 7E-7F, *upper panels*), as we reported previously (Deng et al 2022). Inhibition of Kv7 channels with XE991 (10 μM) further increased AP firing in both genotypes, with significantly higher number of APs still observed in the isolated granule cells of *Fmr1* KO mice (Figures 7E-7F, *lower panels*). These results indicate that intrinsic hyperexcitability of granule cells in *Fmr1* KO mice is largely unrelated to Kv7 channels. Indeed, we have demonstrated that this intrinsic defect is caused by abnormal extrasynaptic GABA_A_ receptor activity in the absence of FMRP (Deng et al 2022). We then performed the same recordings in granule cells with intact dentate circuit, and again observed markedly increased excitability in the *Fmr1* KO, as evident by much larger number of APs fired by KO than WT granule cells (Figures 7G-7H, *upper panels*). In line with our hypothesis, circuit-wide inhibition of Kv7 channels with XE991 abolished the difference in the number of APs between genotypes (Figures 7G-7H, *lower panels*). Together with the observations above (Figure 7F, *lower panel*), these results indicate that XE991 corrected granule cell hyperexcitability in a circuit-dependent manner.

To further clarify this notion, we compared the effects of XE991 between isolated-(Figure 7F) and circuit-intact-granule cells (Figure 7H), and probed the origin of XE991 contribution to granule cell excitability. Specifically, even though the direct action of XE991 on granule cells was to increase excitability in both genotypes (Figure 7—figure supplement 2A), the circuit action of XE991 resulted in suppressing granule cell excitability in *Fmr1* KO mice (-.J ∼6 APs, Figure 7—figure supplement 2B), while having minimal effect in WT mice (changes fluctuating around 0, Figure 7—figure supplement 2B). This is in line with the observations above that XE991 increased sIPSC frequency but not sEPSC frequency, and only in the KO but not in WT mice (Figures 7A-7D). Together with our findings that the interneuron activity is driven in a large part by excitatory input from the MCs, these results indicate that the primary site of action of XE991 is the MCs in *Fmr1* KO mice. Thus, our findings support the notion that circuit-wide inhibition of Kv7 channels boosted up inhibitory drive onto granule cells and restored granule cell excitability in *Fmr1* KO mice by enhancing the local inhibitory circuit function.

### Implications of mossy cell hypo-excitability and circuit E/I imbalance to dentate function

Granule cell excitability and dentate output are not only dependent on basal synaptic inputs (i.e., basal E/I balance, as evident in Figure 7), but more importantly these cellular/circuit functions are controlled by a delicately tuned dynamic E/I balance during circuit activity. By providing the major excitatory drive to local interneurons, MCs play a key role in controlling circuit inhibition onto granule cells via a three-synapse pathway (Granule cells→MCs→Interneurons→Granule cells, Figure 8—figure supplement 3A). Accordingly, we hypothesized that the Kv7-dependent hypo-excitability of MCs in the *Fmr1* KO mice compromises the effectiveness of this critical inhibitory pathway.

To test this possibility, we recorded compound postsynaptic current (cPSC) in granule cells by stimulating PP and holding cells at -45 mV. This holding potential was chosen as it was an intermediate potential between the excitatory and inhibitory reversal potentials ensuring comparable driving force for excitatory and inhibitory conductances. The PP-stimulation-evoked cPSC is the summation of largely overlapping excitatory and inhibitory postsynaptic currents, which shows as an initial down-ward excitatory component and followed by an up-ward inhibitory component (Figure 8—figure supplement 1A). At the end of each recording, the pure EPSC was isolated by adding GABA_A_ receptor blocker gabazine (5 μM) (Figure 8—figure supplement 1B), which then was averaged to create an EPSC template for each cell in order to isolate underlying EPSC and IPSC from the cPSC and calculate E/I ratio (Figure 8—figure supplement 1C-1D). For better comparison among cells, we normalized the cPSC and underlying IPSC to their respective underlying EPSC, which reflects the PP stimulation intensity (Figure 8—figure supplement 1E).

In line with our observations above, loss of FMRP significantly decreased the inhibitory component of cPSC and the underlying IPSC (Figures 8D-8E, *upper panels*), while the excitatory component of cPSC was comparable between genotypes (Figure 8C, *upper panel*). As a result, E/I ratio of the inputs onto granule cells was abnormally increased in *Fmr1* KO mice (Figures 8F-8G, *upper panels*). Moreover, we also observed a wider excitation window (defined as the duration of cPSC excitatory component, Figure 8—figure supplement 1C) in the granule cells of the KO mice (Figure 8H, *upper panel*). In support of our hypothesis, circuit-wide inhibition of Kv7 with XE991 significantly increased the inhibitory component of cPSC and underlying IPSC in *Fmr1* KO (Figures 8D-8E, *lower panels*), but not in WT mice. Consequently, inhibition of Kv7 normalized E/I ratio (Figures 8F-8G, *lower panels*), as well as the excitation window in the granule cells of *Fmr1* KO mice (Figure 8H, *lower panel*). These results indicate that Kv7-dependent hypo-excitability of MCs in *Fmr1* KO mice is a critical defect in the dentate circuit and that circuit-wide inhibition of Kv7 is sufficient to normalize circuit E/I balance.

**Figure 8.**
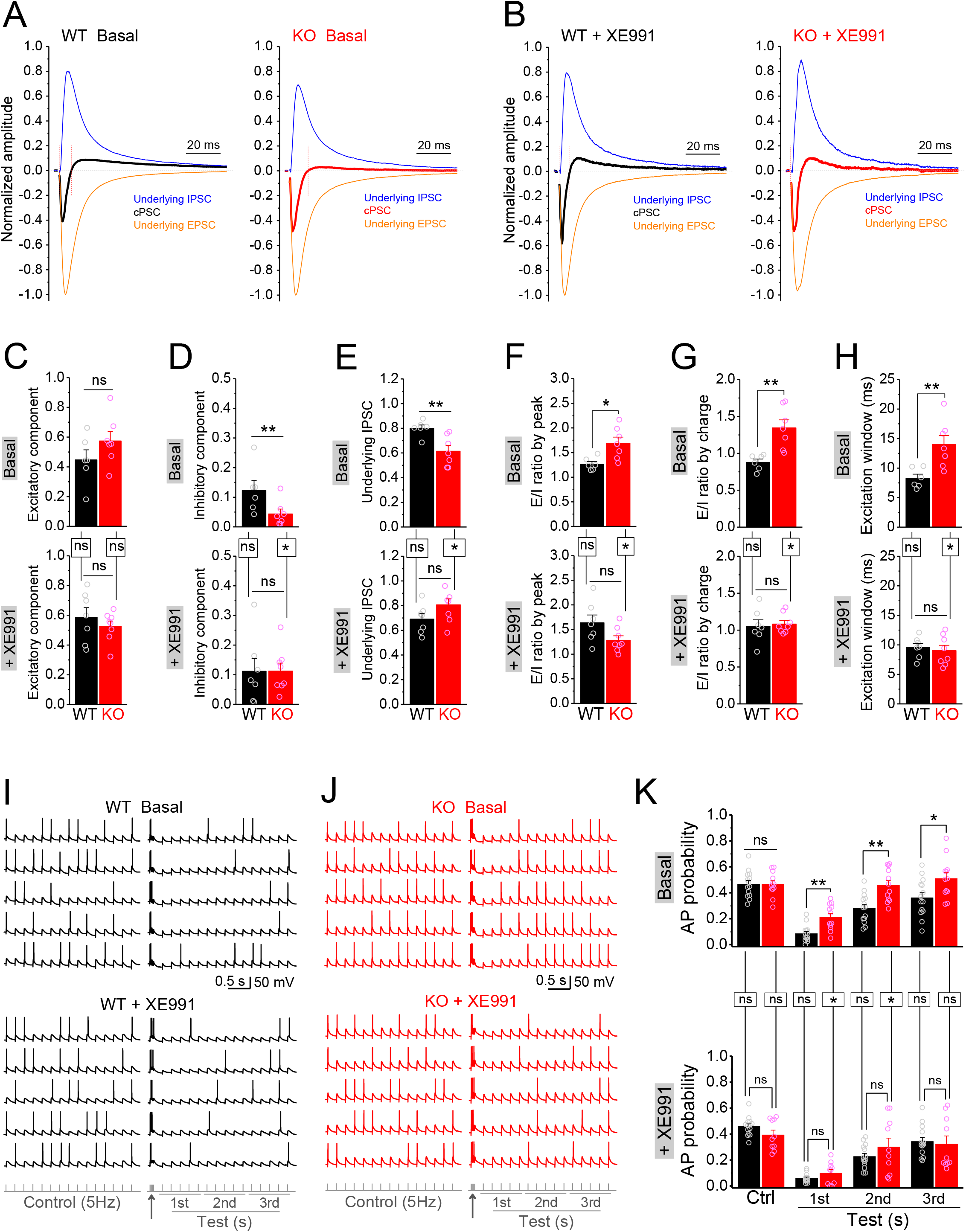
Circuit-wide inhibition of Kv7 channels restored dentate output during theta-gamma coupling stimulation in *Fmr1* KO mice. (A) Sample traces of PP-stimulation evoked compound postsynaptic currents (cPSC) and their respective underlying EPSC and IPSC, in the basal state. For better comparison, the traces were normalized to their own underlying EPSC, which reflects stimulation intensity. Red vertical lines denote the excitation window that was summarized in (H). Stimulation artifacts were removed and baseline before stimulation was shifted to be 0 for presentation purpose. (B) The same as in (A), but in the presence of XE991. (C-E) Summary data of normalized excitatory component (C), inhibitory component (D) and underlying IPSC (E), in the absence (basal, *upper panels*) and in the presence of XE991 (+ XE991, *lower panels*). (F) Summary data of E/I ratio evaluated by the peaks of underlying EPSC and IPSC in the absence (basal, *upper panel*) and in the presence of XE991 (+ XE991, *lower panel*). (G) The same as in (F), but evaluated by the charge transfers of underlying EPSC and IPSC. (H) Summary data of excitation window in the absence (basal, *upper panel*) and in the presence of XE991 (+ XE991, *lower panel*). (I) Sample traces of theta-gamma coupling stimulation-evoked APs in granule cells from WT mice, in the absence of (basal, *upper panel*) or in the presence of XE991 (+XE991, *middle panel*). *Lower panel* showing stimulation protocols: Control, 15 stimuli at 5 Hz; Test, a burst of gamma stimulation (5 stimuli at 50 Hz, *arrow*) 200 ms before 15 stimuli at 5 Hz. AP probability in test train was calculated in 1-second-bin (ie, binned in 1st, 2nd or 3rd second) and plotted in (K). (J) The same as in (I), but for KO mice. (K) Summary data of gamma-suppression of dentate output in response to PP stimulation at theta frequency in the absence of (basal, *upper*) or in the presence of XE991 (+XE991, *lower*). Note loss of FMRP compromised gamma-suppression of AP output in granule cells, and Kv7 blocker XE991 restored the gamma burst-induced suppressive effect on dentate output in *Fmr1* KO mice. *p < 0.05; **p < 0.01; ns, not significant. Horizontal lines (with or without dropdown) denote comparison between genotypes; vertical lines indicate comparison between in the absence and presence of XE991 within genotypes. The statistical data are listed in Supplementary file 1– Supplementary Table 1. Data are mean ± SEM.

Finally, we asked what are the functional consequences of MC hypo-excitability and dentate circuit E/I imbalance in *Fmr1* KO mice? The power of theta-gamma oscillations is particularly high in the dentate gyrus (Csicsvari et al 2003), and plays a critical role in many dentate functions, such as pattern separation (Leutgeb et al 2007), information coding (Mizuseki et al 2009, Pernia-Andrade & Jonas 2014), and thus learning/memory (Bott et al 2016, Lisman & Jensen 2013, Neves et al 2022). In order to evaluate the contribution of MC defects in *Fmr1* KO mice to this critical dentate function, we used a protocol of double-oscillation interplay at the single-cell level (Hasselmo, Giocomo & Zilli 2007, Mircheva, Peralta & Toth 2019). In this paradigm, gamma (∼50Hz) frequency stimulation of PP suppresses granule cell output driven by PP stimulation at the theta (∼5Hz) frequency range (gamma-suppression, for brevity). The experiments included two stimulation protocols: a control protocol at 5 Hz (theta stimulation, whose intensity was adjusted to achieve AP probability of ∼0.5 and the intensity was then kept in the following test protocol for the same cell); and a test protocol with a gamma burst (5 stimuli at 50 Hz) included 200 ms before the 5 Hz theta train. Theta stimulation of the PP evoked a steady baseline firing in granule cells (Figures 8I-8K, *upper panels*). In line with previous studies (Hasselmo, Giocomo & Zilli 2007, Mircheva, Peralta & Toth 2019), a preceding gamma stimulation suppressed granule cell output in response to theta train in both genotypes (Figures 8I-8K, *upper panels*). However, this gamma-suppression was much less efficient in the KO than in WT mice, as evident by the significant higher AP probability in KO mice (Figures 8I-8K, *upper panels*). Considering that granule cells are the only output neurons in the dentate gyrus and thus integrate all dentate circuit operations, these results indicate that loss of FMRP causes abnormal dentate information processing, leading to excessive dentate output.

The dentate GABAergic system plays a critical role in the above gamma-suppression of granule cell output (Mircheva, Peralta & Toth 2019). Based on our findings, we hypothesized that the compromised dentate inhibitory pathway in the *Fmr1* KO mice weakens gamma-suppression through enhancing the EPSP summation in granule cells. The EPSP measured in granule cells in response to the PP stimulation integrates both excitatory and inhibitory synaptic inputs onto granule cells, including the direct synaptic input from the PP and all the PP stimulation-associated feedforward and feedback synaptic inputs. In other words, the EPSP in granule cells integrates all dentate circuit “operations”. Mechanistically, once the integrated EPSP reaches the threshold level, it triggers an AP, which makes the EPSP amplitude a direct predictor of AP firing. We found that theta stimulation evoked a steady EPSP (Figure 8—figure supplement 2A-2B), and a preceding gamma burst markedly suppressed the EPSP in WT mice (Figure 8—figure supplement 2A-2B). In line with our results above, this suppression effect was significantly weaker in KO mice (Figure 8—figure supplement 2A-2B), suggesting that the compromised dentate inhibitory pathway weakens gamma-suppression through enhancing the EPSP summation in the KO mice (Figure 8—figure supplement 3B). If this is the case, our results predict that circuit-wide inhibition of Kv7 channels should dampen EPSP summation and restore the gamma-suppression. Indeed, we found that Kv7 blocker XE991 reduced the EPSP amplitude in KO mice to the WT level (Figure 8—figure supplement 2C-2D) and, most importantly, it normalized gamma-suppression of granule cell output during theta activity in *Fmr1* KO mice (Figures 8J-8L and Figure 8—figure supplement 3C).

Collectively, these findings demonstrates that MCs hypo-excitability in *Fmr1* KO mice results in circuit E/I imbalance that compromises the dentate information processing functions, and that circuit-based interventions, such as inhibition of Kv7, could be a potential therapeutic strategy to re-normalize the circuit E/I balance and ameliorate dentate dysfunction in FXS.

## Discussion

Dentate gyrus, the information gateway to the hippocampus, receives and processes the vast majority of cortical-hippocampal inputs. It carries out numerous information-processing tasks to form unique memories, and conveys the information to the CA3 region. The precisely controlled E/I balance in the dentate circuit is critical to many fundamental computations that are believed to underlie learning and memory (Amaral, Scharfman & Lavenex 2007). The dentate gyrus output is controlled by the local networks whose activity is determined not only by the entorhinal cortical inputs and local interneuron activity, but also by another type of local excitatory neurons, the MCs (Scharfman 2016, Scharfman 2018). These cells has long been considered “enigmatic” due to their complex and incompletely understood roles in the circuit, yet recently the critical roles of MCs in dentate computations and spatial memory have emerged (Amaral, Scharfman & Lavenex 2007, Berron et al 2016, Botterill et al 2021b, Fredes & Shigemoto 2021, Scharfman 2016). Moreover, MC defects were also found to play major roles in epilepsy and many neurodevelopmental disorders (Pelkey et al 2017, Scharfman 2016, Scharfman 2018, Sloviter et al 2012). Whether MCs defects are present and contribute to hippocampal dysfunction in FXS has remained largely unknown. Here we demonstrate that loss of FMRP causes profound MC hypo-excitability, an unusual defect which is opposite to cellular hyperexcitability reported in all other excitatory neurons in FXS mouse model, including dentate granule cells (Deng et al 2022). Since MC hypo-excitability was observed both in the synaptically isolated MCs and with the intact circuit, this defect has a cell-autonomous origin and the adaptive circuit changes were insufficient to compensate for it.

In pursuing the mechanism of this defect, we discovered that it is caused by abnormally elevated Kv7 function in the absence of FMRP. While Kv7 contribution to input resistance is negligible at the resting potential of MCs (below -60 mV) (Delmas & Brown 2005, Jones, Gamper & Gao 2021, Liu, Bian & Wang 2021), once the membrane potential rises to a near-threshold level (-45 mV), the excessive activity of Kv7 channels act to reduce input resistance and suppress AP initiation thus reducing MC excitability in *Fmr1* KO mice. How does loss of FMRP cause an increase in Kv7 function? Both KCNQ2 and KCNQ3 have been identified as FMRP translational targets (Darnell et al 2011). Our western blot analysis showed that the expression levels of KCNQ2 and KCNQ3 in KO mice were not altered in the whole brain lysate in general, nor in the dentate gyrus region, specifically. In addition to translational regulation, FMRP is also known to regulate activity of several K^+^ channels, including Slack, BK, SK and Kv1.2 via protein-protein interactions (Deng & Klyachko 2021). However, in all of these cases, FMRP loss decreases rather than increases K^+^ channel activity, making loss of protein-protein interactions an unlikely mechanism to explain increased Kv7 function. One alternative possibility is that loss of FMRP affects intracellular second-messenger signaling pathways that may regulate Kv7 activity (Haick et al 2017). For example, activation of PKC suppresses Kv7 function (Delmas & Brown 2005, Haick et al 2017) and PKCε expression was found to be reduced in hippocampus of *Fmr1* KO mice (Marsillo et al 2021), supporting a possibility that the increased Kv7 function in KO MCs may be associated with abnormal PKC signaling. However, further studies will be needed to elucidate the precise mechanism responsible for the increased Kv7 function in *Fmr1* KO mice.

Our analyses show that while the MC excitability dysfunction is unusual in being opposite in direction to other excitatory neurons, it nevertheless leads to abnormally increased granule cell output (i.e., dentate output) in *Fmr1* KO mice. This counterintuitive effect arises because in addition to directly exciting granule cells, MCs act via local inhibitory networks to indirectly suppress granule cell activity. Therefore, the net effect of MC activity on granule cell output is orchestrated via a dynamically tuned E/I balance of direct excitation and indirect inhibition. In their control of inhibition, the MCs relay and expand granule cell-derived excitatory drive onto interneurons, which then convert this drive into stronger feed-back inhibition onto granule cells (Botterill et al 2021a, Hashimotodani et al 2017, Houser et al 2021, Yeh et al 2018). Notably, the resulting E/I balance driving granule cell output is not static; rather, during network activity, short-term plasticity actively modulates outputs from both excitatory and inhibitory synapses to dynamically fine-tune the E/I drives and optimize neural computations (Bartley & Dobrunz 2015, Bhatia, Moza & Bhalla 2021, Grangeray-Vilmint et al 2018). Consequently, the MC hypo-excitability in *Fmr1* KO mice effectively reduces excitatory drive onto local interneurons, which in turn reduces local inhibitory drive to granule cells, ultimately leading to abnormal dentate information processing and exaggerated granule cell output. Given that granule cells normally fire sparsely during information processing, this increased AP output in granule cells is reminiscent of abnormal CA3-to-CA1 information transmission in the *Fmr1* KO mice, in which a large amount of ‘‘noise’’ that is normally filtered out during synaptic processing is being transmitted indiscriminately due to FMRP loss (Deng et al 2013).

We note the discrepancy in the changes of inhibitory drives among different cell types, i.e., sIPSC frequency was increased in MCs, showed no change in granule cells and a decrease in interneurons, which cannot be explained by a simple change in excitability of any one cell type in dentate gyrus. Rather, these distinct changes might indicate a compensatory circuit reorganization of the interneuron axonal terminals onto these three targets in the absence of FMRP. Similarly, the observed alterations in excitatory drive onto interneurons, including both mEPSC and sEPSC frequencies, suggest changes in the excitatory synapse number and/or function. Together with alterations in inhibitory drives, these changes may reflect compensatory reorganization of both excitatory and inhibitory connections, including mossy cell synapses. Indeed, FMRP plays a crucial role in synaptogenesis and synaptic refinement (Booker et al 2019, Doll & Broadie 2014, Kennedy, Rinker & Broadie 2020, Sears & Broadie 2018), and loss of FMRP is well known to cause changes in synaptic connectivity (Gibson et al 2008, Kennedy, Rinker & Broadie 2020, Patel et al 2013, Patel et al 2014, Tessier & Broadie 2009). Such circuit reorganization can explain the balanced E/I drive onto granule cells in *Fmr1* KO mice we observed in the basal state, which can result from reorganization of inhibitory axonal terminals in favor of increased contacts onto granule cells and MCs, and reduced contacts onto other interneurons, as a compensation for the decreased MC-derived excitatory drive. In line with this notion, it has been reported that dentate circuit reorganization occurs after status epilepticus, which can increase inhibitory drive onto granule cells and function as a homeostatic compensation (Butler, Westbrook & Schnell 2022). Indeed, many of the cellular and synaptic changes in the *Fmr1* KO mice are antagonistic and mitigating circuit dysfunction, and thus compensate for the primary defects (Domanski et al 2019). However, even though the homeostatic circuit compensation is able to maintain the basal E/I input balance in granule cells of *Fmr1* KO mice, this compensation was insufficient to maintain dynamic E/I balance during active states, such as during network activity evoked by PP stimulation. Consequently, the dentate output was exaggerated during the physiologically-relevant theta-gamma coupling activity in *Fmr1* KO mice. Indeed, as discussed above, E/I balance is dynamically tuned to rapidly adjust during circuit computations (Bartley & Dobrunz 2015, Bhatia, Moza & Bhalla 2021, Grangeray-Vilmint et al 2018), which is thought to represent a fundamental mechanism underlying efficient neural coding (Zhou & Yu 2018). Considering that the theta-gamma oscillation coupling plays a critical role in dentate information processing (Bott et al 2016, Leutgeb et al 2007, Lisman & Jensen 2013, Mizuseki et al 2009, Neves et al 2022, Pernia-Andrade & Jonas 2014), our findings indicate that MC-driven E/I imbalance is a critical defect contributing to dentate dysfunction in FXS. Moreover, by projecting axons both ipsi-and contra-laterally along the longitudinal axis of the dentate gyrus, MCs are uniquely positioned to influence dentate activity widely across lamellae (Jinde, Zsiros & Nakazawa 2013, Scharfman 2016, Scharfman 2018). This wide-reaching control of dentate activity makes MC hypo-excitability in *Fmr1* KO mice a core defect in the dysregulation of circuit E/I dynamics and information processing.

Most critical discovery in this regard is our finding that circuit-wide inhibition of Kv7 was sufficient to re-normalize E/I balance in the dentate circuit despite the irreconcilable hypo-excitable state of MCs and hyperexcitable state of granule cells in the *Fmr1* KO mice. Specifically, XE991 corrected MC hypo-excitability via direct effect on Kv7 channels in MCs and normalized granule cell excitability through dentate three-synapse feedback pathway in *Fmr1* KO mice. Moreover, inhibition of Kv7 channels was also sufficient to restore the dentate output during the physiologically-relevant theta-gamma coupling activity in KO mice. Although inhibition of K^+^ channels is typically expected to increase circuit excitability, the key role of MCs as the main excitatory drive for this three-synapse feed-back inhibition resulted in the overall suppression of circuit hyperexcitability in the dentate circuit of *Fmr1* KO mice. This effect arises because the excessive Kv7 function is present selectively in MCs of *Fmr1* KO mice. As a result,inhibition of Kv7 had a stronger effect on enhancing MC-dependent feedback inhibition onto granule cells than directly increasing granule cell excitability in *Fmr1* KO mice. Notably, the applicability of a circuit-wide approach as a potential treatment *in vivo* will require extensive future behavioral analyses, which are beyond the scope of the current study.

Taken together, these findings provide a proof-of-principle demonstration that a circuit-based intervention can normalize dynamic E/I balance and restore dentate circuit output *in vitro*. Thus, circuit-based interventions can represent a potential therapeutic strategy to correct dentate dysfunction in this disorder.

## Materials and methods

### KEY RESOURCES TABLE

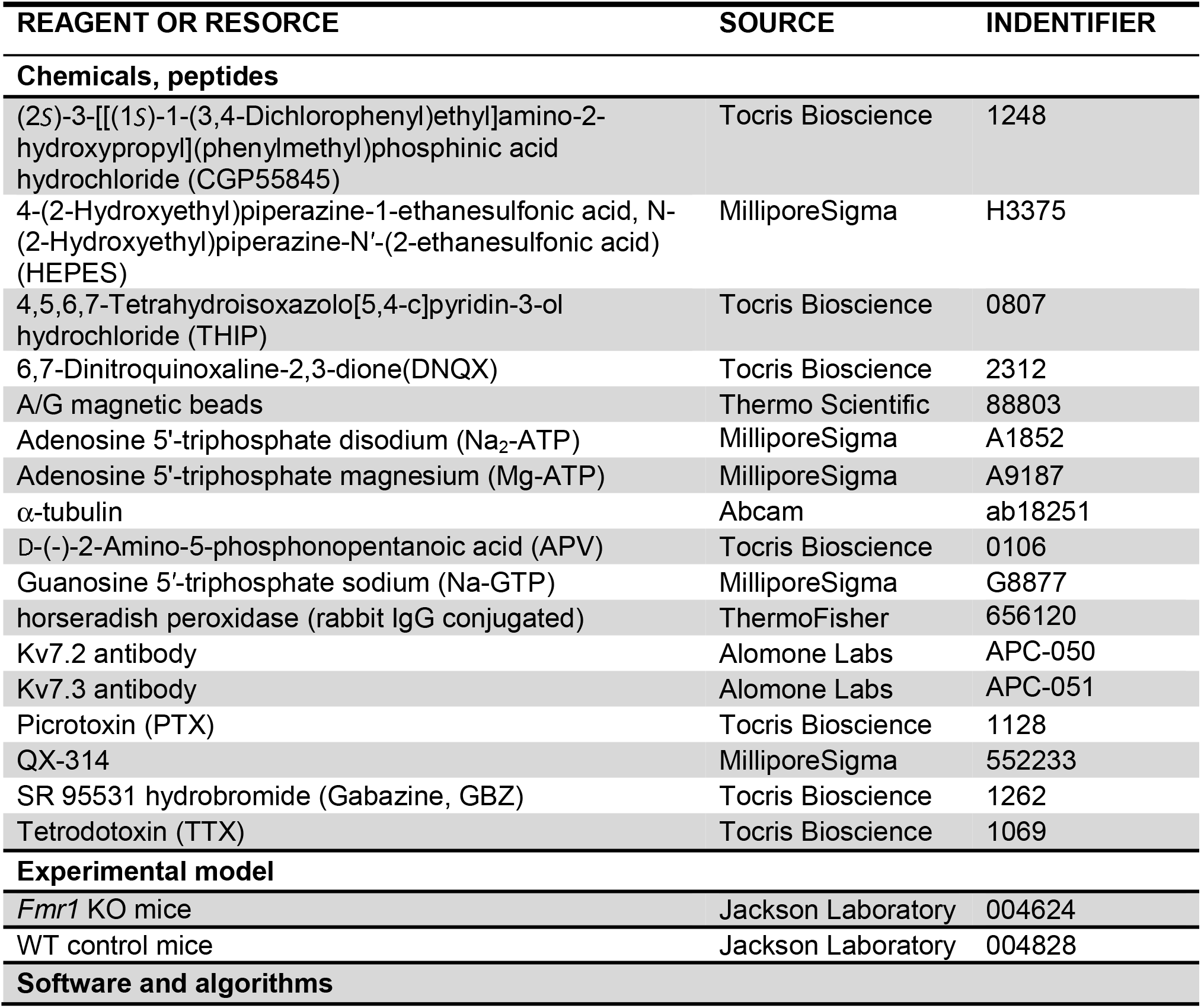
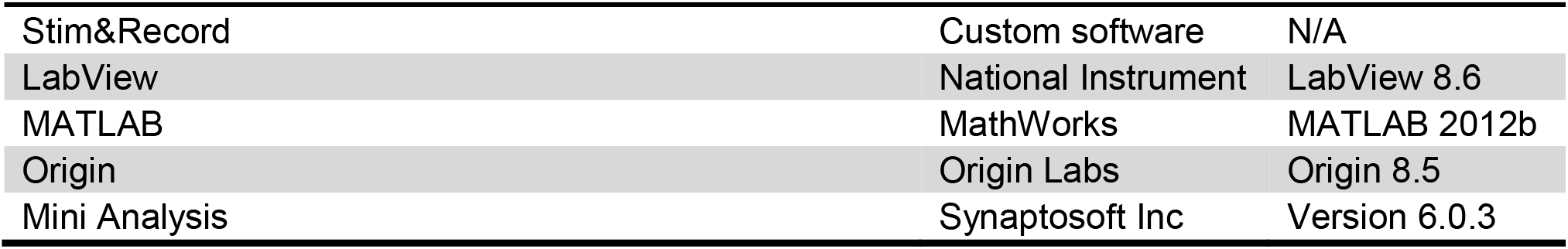

#### Animals and slice preparation

*Fmr1* KO (FVB.129P2-Pde6b^+^ Tyr^c-ch^ Fmr1^tm1Cgr^/J; stock #004624) and WT control mice (FVB.129P2-Pde6b^+^Tyr^c-ch^/AntJ; stock #004828) were obtained from The Jackson Laboratory. Slices were prepared as previously described (Deng et al 2022). In brief, male 21―23-day-old mice were used. After being deeply anesthetized with CO_2_, mice were decapitated and their brains were dissected out in ice-cold saline containing the following (in mm): 130 NaCl, 24 NaHCO_3_, 3.5 KCl, 1.25 NaH_2_PO_4_, 0.5 CaCl_2_, 5.0 MgCl_2_, and 10 glucose, pH 7.4 (saturated with 95% O_2_ and 5% CO_2_). Horizontal hippocampal slices (350 μm) were cut using a vibrating microtome (Leica VT1100S) (Deng et al 2022). Slices were initially incubated in the above solution at 35°C for 1 h for recovery and then kept at room temperature (∼23°C) until use. All animal procedures were in compliance with the US National Institutes of Health Guide for the Care and Use of Laboratory Animals, and conformed to Washington University Animal Studies Committee guidelines.

#### Action potential recording

AP recordings using a Multiclamp 700B amplifier (Molecular Devices) were made from mossy cells, interneurons or granule cells with infrared video microscopy and differential interference contrast optics (Olympus BX51WI). All of the recordings were conducted at near-physiological temperature (33–34°C). The recording electrodes were filled with the following in the present study (unless stated otherwise) (in mM): 130 K-gluconate, 10 KCl, 0.1 EGTA, 2 MgCl_2_, 2 ATPNa_2_, 0.4 GTPNa, and 10 HEPES (pH 7.3). The extracellular solution contained (in mM): 125 NaCl, 24 NaHCO_3_, 3.5 KCl, 1.25 NaH_2_PO_4_, 2 CaCl_2_, 1 MgCl_2_, and 10 glucose, pH 7.4 (saturated with 95% O_2_ and 5% CO_2_).

Mossy cells were identified by their location (hilus), characteristic morphology (size and shape) and electrophysiological properties, including RMP (∼-65 mV), cell capacitance (>100 pF), input resistance (∼150 MΩ at RMP), sEPSCs with high frequency and large amplitude (>60 pA), AP with relatively small afterhyperpolarization (∼-6 mV) (Bui et al 2018, Chancey et al 2014, Jinde et al 2012, Scharfman & Myers 2013, Wang et al 2021). For determination of excitability, mossy cell membrane potential was set to given potentials (-64 to -55 mV with 1-mV step) through amplifier’s function of automatic current injection. Number of APs within 20 s was averaged from 4-5 trials for each cell. For measurements of AP parameters, we employed a ramp current injection (0.15 pA/ms) (Deng et al 2019, Deng et al 2022) with a hyperpolarizing onset to ensure maximal Na^+^ channel availability before the 1st AP. All AP parameters (except number of APs) were determined only from the first APs of ramp-evoked AP trains to avoid the influence from cumulatively inactivating voltage-gated channels. AP threshold was defined as the voltage at the voltage trace turning point, corresponding to the first peak of 3rd order derivative of AP trace (Deng et al 2022). The rheobase was defined as the ramp current amplitude at the time point of threshold; rheobase charge transfer was determined by the integration of time and input current. The maximum rising rate was determined as the peak of the 1st order derivative of AP trace. AP duration was measured at the level of AP trace crossing -10 mV. The AP amplitude was measured from the threshold point to the AP peak because the membrane potential was continuously ramp-up with time and the threshold point is the most reliable point before an AP in this setting. The rise and fall time was the interval of 10–90% amplitude during AP upstroke and down-stroke, respectively.

Hilar interneurons were easily distinguished from mossy cells according to the differences in morphology and electrophysiological properties (Lubke et al 1998) [relative to mossy cells, interneurons have more depolarizing RMP (∼-55 mV), smaller capacitance (<60 pF), higher input resistance (usually >300 MΩ at RMP), and larger afterhyperpolarization (lower than -15 mV)]. AP firing pattern in response to a 100 pA depolarizing step current for 600 ms was used to classify interneurons into 3 types (Figure 5—figure supplement 1A).

Granule cells were identified by the location (granule cell layer) and characteristic electrophysiological properties (relative to interneurons) (Deng et al 2022): very hyperpolarizing resting membrane potential (∼ -80mV), very low sEPSC frequency (<20 events/min), and smaller capacitance (∼20 pF). Cells that could not be definitively classified into the three categories above were not included in further analyses. To avoid recording from newly generated immature granule cells, we used cells located at the outer regions of the granule cell layer (Deng et al 2022). For step-current-evoked AP output in granule cells, we used multistep-current (50–75 pA with an increase of 5 pA/step for 600 ms) to evoke APs from granule cells. The RMP was set to -80 mV by constant current injection (if necessary) for better comparison among cells. Number of APs was averaged over 5-8 trials in each cell. For PP stimulation-evoked AP in granule cells, the stimulation protocol consisted of a train of 5 Hz; the stimulation electrodes positioned in the middle molecular layer of dentate gyrus to stimulate medial PP. The RMP of granule cells was set to -70 mV for facilitating AP firing and the stimulation intensity was adjusted so that the AP probability was ∼0.5 (under 5 Hz stimulation). A burst of gamma stimulation (5 stimuli at 50 Hz, 200 ms before the first stimulus of 5 Hz train) was used to evaluate gamma stimulation-induced suppression of granule cell output in response to PP theta stimulation. Granule cell output was expressed as the AP probability. When PP stimulation failed to evoke APs, we measured the amplitude of EPSPs, which was defined as the voltage differences between individual EPSP peaks and the initial baseline before gamma-stimulation (set to be -70 mV). For better comparison, the EPSP amplitudes were normalized to their own controls. AP probability was calculated from at least 10 trials in each cell.

#### Measurement of resting membrane potential, capacitance and input resistance

Resting membrane potential (RMP) was measured immediately after whole-cell formation. Cell capacitance is determined by the amplifier’s auto whole-cell compensation function with slightly manual adjustment to optimize the measurement if needed. For input resistance, assessment was performed using the canonical method, in which the voltage response to a negative current injection (−60 pA for 500 ms) at RMP was used to calculate input resistance. For measurement of input resistance around threshold levels, we pharmacologically isolated MCs using blockers against both glutamate and GABA receptors (in μM, 10 NMDA, 50 APV, 10 MPEP, 5 gabazine and 2 CGP55845) to isolate mossy cells from the dentate circuit. TTX (1 μM) and CdCl_2_ (10 μM) were also used to block Na^+^ and Ca^2+^ channels, respectively. Under these conditions, a positive current injection (+60 pA for 500 ms) at -45 mV (set by constant current injection) was used to evoke voltage response, which then used to calculate input resistance.

#### Determination of changes in holding current and membrane potential

Holding current were recorded by holding cells at -45 mV in pharmacologically isolated MCs, and in the presence of TTX (1 μM) and 10 μM CdCl_2_. Changes in holding current were the differences in holding current before and during XE991 (10 μM). For determining changes in membrane potentials in response to Kv7 inhibition, membrane potentials were initially set at -45 mV (by constant current injection). Under these conditions, the differences in membrane potentials before and during XE991 (10 μM) were calculated as the changes in membrane potential.

#### Kv7 current recording

We used a depolarizing voltage ramp (-95 to +5 mV, 0.02 mV/ms) to evoke Kv7 current from MCs, using the same internal and external solutions as those in holding current recordings. Cell capacitance was compensated. Series resistance compensation was enabled with 80–90% correction and 16 μs lag. Kv7 current was isolated by subtracting current in 10 μM XE991 from that before XE991. The I-V curves were constructed from ramp-evoked Kv7 currents every 5 mV and normalized to respective cell capacitances (mean current value over 0.01 mV intervals from averages of four to five trials for each cell to approximate quasi-steady-state current).

#### Recordings of spontaneous and miniature postsynaptic currents

Spontaneous excitatory postsynaptic currents (sEPSCs) were recorded from hilar mossy cells (holding at -65 mV) and interneurons (at -60 mV). The pipette solution was the same as that used in AP recording, except that QX-314 (1 mM) was included in the pipette solution to block possible action current. The bath solution was supplemented with gabazine (5 μM) to block GABA_A_R responses. The solutions used to recording of miniature excitatory postsynaptic currents (mEPSCs) were the same as those for sEPSCs, except that tetrodotoxin (TTX, 1 μM) was included in the bath solution to block action potential-dependent responses.

For recording of spontaneous inhibitory postsynaptic currents (sIPSCs), the recording pipette solution contained (in mM): 130 CsCl, 2 MgCl2, 4 Mg-ATP, 0.3 Na-GTP, 10 HEPES, and 0.1 EGTA (pH 7.3). The bath solution was supplemented with APV (50 μM) and DNQX (10 μM) to block responses of ionotropic glutamate receptors. Note that IPSC is down-going signal in this high chloride internal solution. For miniature inhibitory postsynaptic currents (mIPSCs) recording, TTX was added in the bath solution.

For simultaneous recording of sEPSCs and sIPSCs from granule cells, the recording pipette solution contained (in mM): 135 K-gluconate, 2 MgCl_2_, 0.1 CaCl_2_, 2 MgATP, 0.3 NaGTP, 4 Na_2_-phosphocreatine, 0.2 EGTA and 10 HEPES (pH 7.3). External solution was the same as AP recordings (without any blockers, unless stated otherwise). The membrane potential was held at -40 mV. Under this condition, EPSC is down-going signal (inward current) and IPSC up-going signal (outward current). The events detection threshold was 10 pA for sEPSC and sIPSC detection; the automatic detection was visually verified.

#### Recording of compound postsynaptic current and isolation of underlying EPSC and IPSC

The compound postsynaptic current (cPSC) was recorded from granule cells by stimulation medial PP at 0.2 Hz and using the same pipette solution as that of simultaneous recording of sEPSCs and sIPSCs, except that QX314 (1 mM) was included to block action current. Granule cells were held at -45 mV, which is an intermediate potential between the excitatory and inhibitory reversal potentials ensuring comparable driving force for excitatory and inhibitory conductances. The PP-stimulation-evoked cPSC had an initial down-ward excitatory component and followed by an up-ward inhibitory component (Figure 8—figure supplement 1A). The cPSC excitation window was defined as the full duration of excitatory component (Figure 8—figure supplement 1C). At the end of each recording, the pure EPSC (Figure 8—figure supplement 1B) was recorded by adding GABA_A_ blocker gabazine (5 μM) and keeping the same stimulation intensity, which was used to create an EPSC template (average of at least 20 uncontaminated EPSCs) for each cell. The EPSC template was then repeatedly scaled to each data point of cPSC “approximating segment” to obtain a set of scaled EPSCs (Figure 8—figure supplement 1C). All scaled EPSCs were then averaged to approximate an underlying EPSC for a given cPSC (Figure 8—figure supplement 1C). The “approximating segment” of cPSC was defined as 25–65% height of cPSC excitatory component, but not beyond 2.5 ms after stimulation. The underlying IPSC was isolated by subtracting the underlying EPSC from corresponding cPSC (Figure 8—figure supplement 1D). For better comparison, we normalized the cPSC and underlying IPSC to their respective underlying EPSC, which reflects the PP stimulation intensity (Figure 8—figure supplement 1E). The underlying EPSC and IPSC were then used to estimate E/I ratio by their peak amplitudes or charge transfer (amplitude-time integration within 100 ms). Data for each cell were averages from 20–25 uncontaminated cPSC events.

#### Western blotting

Whole brains or dentate gyrus regions (isolated from the brain slices) were lysed in 2% sodium dodecyl sulfonate (SDS) with protease and phosphatase inhibitors (Roche Applied Sciences) and manually homogenized. Protein concentration was determined by DC protein assay (Bio-Rad Laboratories) against bovine serum albumin standards. 50 μg total protein was run on NuPage 4-12% Bis-Tris polyacrylamide gels (Life Technologies) and transferred to nitrocellulose membranes. Membranes were blotted with antibodies directed against the following proteins: α-tubulin, Kv7.2, Kv7.3, rabbit IgG conjugated to horseradish peroxidase. Blots were developed with SuperSignal West Dura (ThermoFisher) and imaged with a ChemiDoc MP imaging system (Bio-Rad Laboratories).

#### Data and code availability

This paper does not report standardized data types. The data generated and analyzed in this study are included in the manuscript, figures and supplemental information. This study did not generate new or unique code, reagents or other materials.

#### Statistical analysis

The data were analyzed in MatLab, except for the postsynaptic currents (sEPSC, mEPSC, sIPSC and mIPSC) that were analyzed by MiniAnalysis. All figures were made in Origin or MatLab. Data are presented as mean ± SEM. Student’s t test, one-way ANOVA, Kolmogorov–Smirnov (K-S) test or Chi-square test were used for statistical analysis as appropriate. Significance was set as p < 0.05. The n was number of cells tested in electrophysiological experiments, which was from at least 3 different mice for each condition. The N in Western blotting was number of animals used. All statistical values can be found in Supplementary file 1– Supplementary Table 1.

## Supplementary files

Supplemental information can be found at the end of the manuscript, including 8 Supplementary Figures and 1 Supplementary Table.

## Author contributions

PYD and VAK conceived and designed the experiments. PYD performed all electrophysiological studies and analyzed the data. AK and VC designed biochemical experiments. AK and PYD performed these experiments. PYD and VAK wrote the initial manuscript, and all authors edited and approved the final version.

## Declaration of interests

The authors declare no competing interests.

## Acknowledgements

This work was supported in part by NIH grant R35 NS111596 to VAK, and R01 NS111719 and R35 NS122260 to VC. The schematic illustration of Figure 1A was created by Yuhan Deng.

---

**Figure 3—figure supplement 1.**
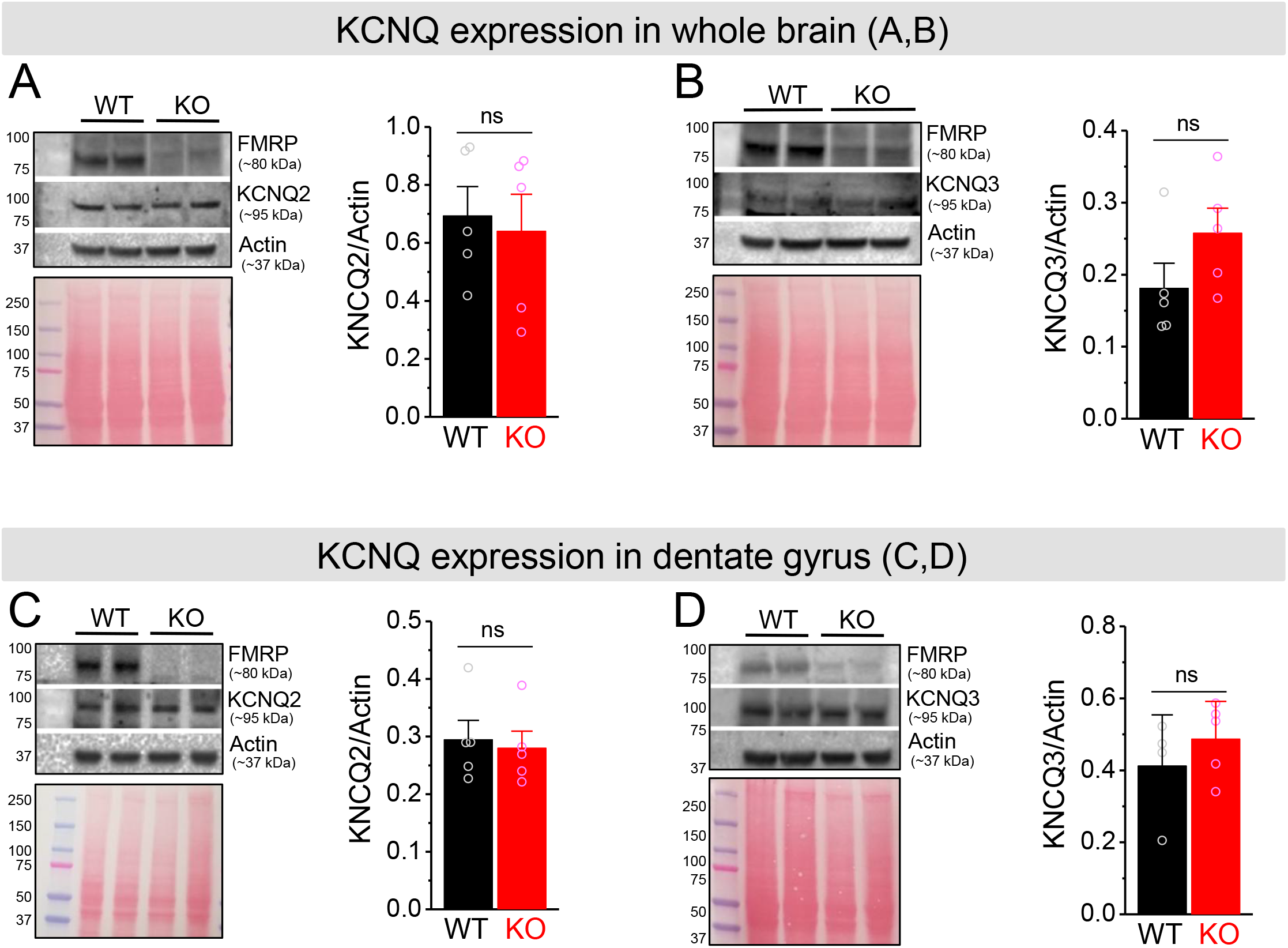
No changes in KCNQ2 and KCNQ3 expression in *Fmr1* KO mice. **a**-**b** Western blot analysis of KCNQ2 (**a**) and KCNQ3 (**b**) in the whole brain lysate. Ponceau staining was used as a loading control for the lysate (*lower panel*). **c**-**d** The same as (**a**-**b**), but for dentate gyrus lysate. ns, not significant. The statistical data are listed in Supplementary file 1–Supplementary Table 1. Data are mean ± SEM.

**Figure 4—figure supplement 1.**
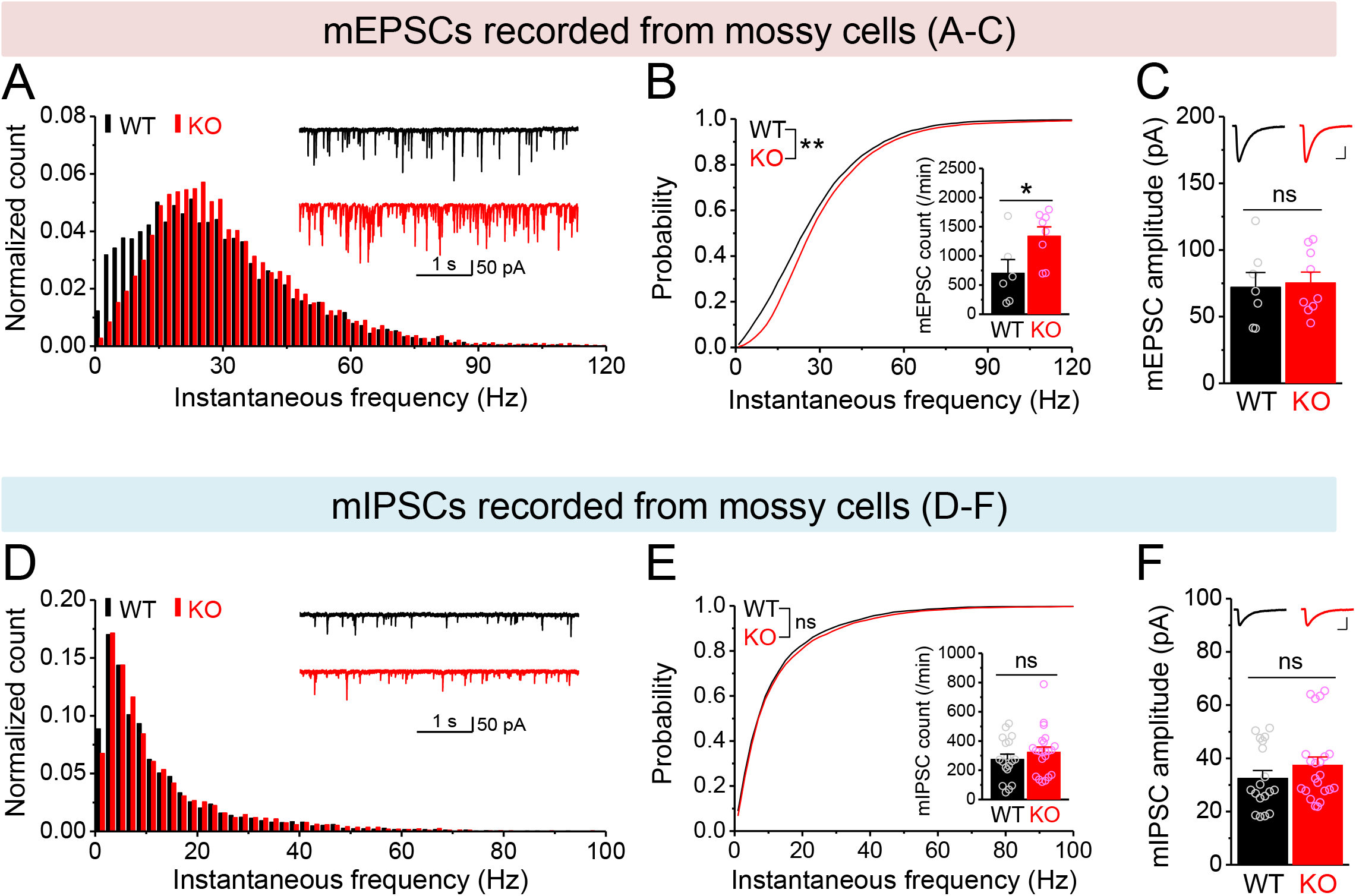
Changes in miniature synaptic inputs onto mossy cells in *Fmr1* KO mice. (A) Distribution of mEPSC instantaneous frequency in MCs. A bin size of 2 Hz was used to calculate mEPSC frequency distribution from a 30-s-long trace per cell. The number of mEPSCs within each bin was normalized to the total number of the respective cells for pooling the data from all cells. Note that mEPSC events in KO mice had a shift toward high frequency. *Insert*, sample traces of mEPSCs for WT (black) and KO (red) mice. (B) Cumulative probability of mEPSC instantaneous frequency in MCs. *bar graph*, number of mEPSCs per minutes. Note that both cumulative probability and number of mEPSCs reveal increased excitatory drive onto MCs. (C) Summary data for mEPSC amplitude. *Insert,* sample mEPSC events for WT (black) and KO (red) MCs. Scale: 5 ms (horizontal) and 25 pA (vertical). (D-F) mIPSCs recorded from KO and WT MCs, aligned in the same way as in (A-C), respectively. *p < 0.05; **p < 0.01. The statistical data are listed in Supplementary file 1–Supplementary Table 1. Data are mean ± SEM.

**Figure 5—figure supplement 1.**
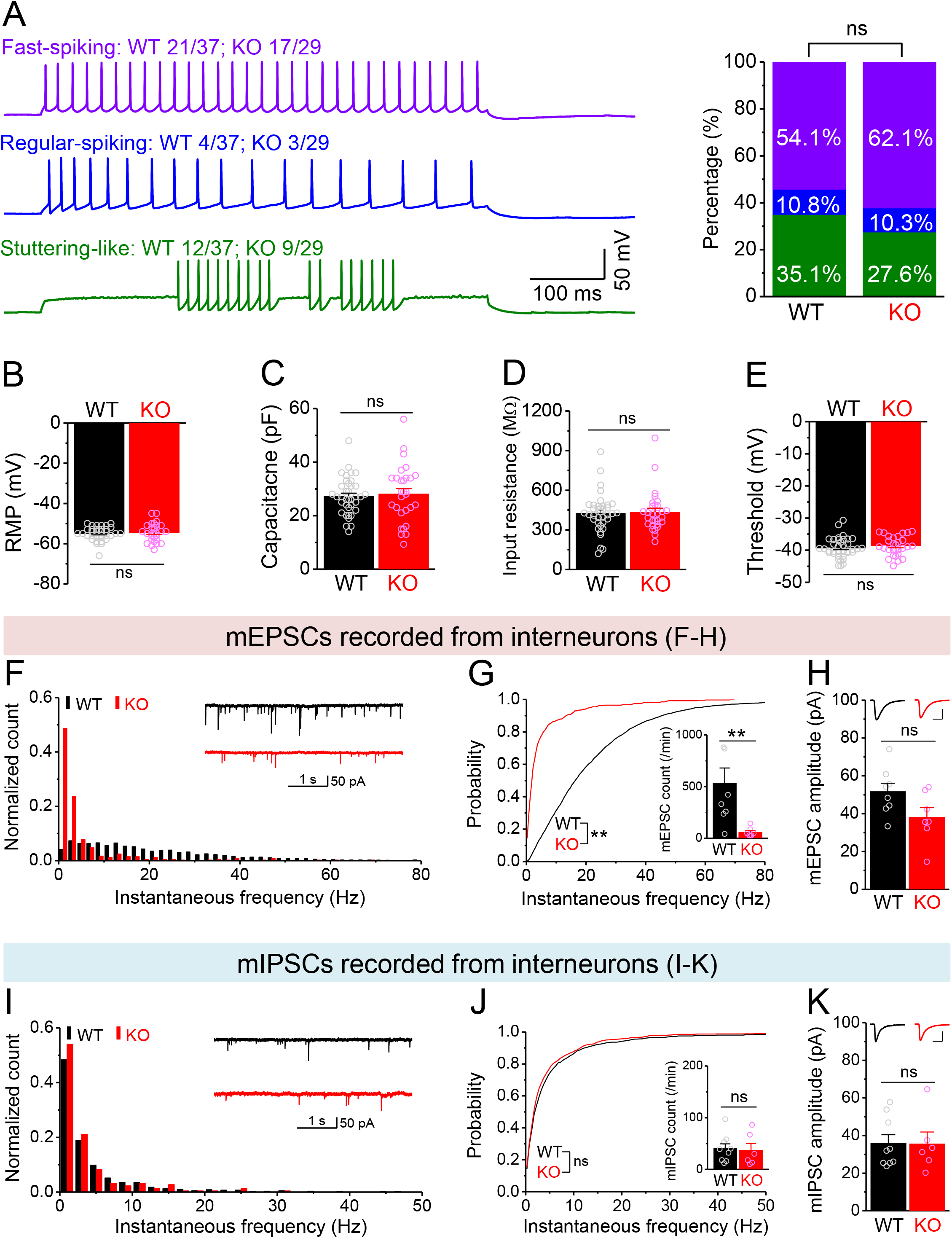
Changes in miniature synaptic inputs onto hilar interneurons in *Fmr1* KO mice. (A) Sample traces (*Left*) of 3 types of interneurons classified by firing pattern. *Upper trace*, fast-spiking interneurons with high-frequency non-adapting firing (WT 20/37 and KO 18/29, where numerator denotes the number of cells for a given type and denominator is the total number of tested cells for a given genotype); *Middle trace*, regular-spiking interneurons with slower and adapting firing (WT 4/37, KO 3/29); *Lower trace*, stuttering-like interneurons with high-frequency irregular bursting firing (WT 13/37, KO 8/29). Stack bar graph (*Right*) shows the percentage of 3 interneuron types (color-coded as the traces in the *Left panel*). No significant differences in the ratios of 3 interneuron types between WT and KO mice. (B-E) We pooled RMP (B), capacitance (C), input resistance (D) and threshold (E) from 3 types of interneurons due to no significant differences observed among types. (F) Distribution of mEPSC instantaneous frequency in interneurons. A bin size of 2 Hz was used to calculate mEPSC frequency distribution from a 30-s-long trace per cell. The number of mPSCs within each bin was normalized to the total number of the respective cells for pooling the data from all cells. Note that mEPSC events in KO mice had a shift toward low frequency. *Insert*, sample traces of mEPSCs for WT (black) and KO (red) mice. (G) Cumulative probability of mEPSC instantaneous frequency in interneurons. *Bar graph* shows number of mEPSCs per minutes. Note that both cumulative probability and number of mEPSCs reveal decreased excitatory drive onto interneurons. (H) Summary data for mEPSC amplitude. *Insert,* sample mEPSC events for WT (black) and KO (red) MCs. Scale: 5 ms (horizontal) and 25 pA (vertical). (I-K) mIPSCs recorded from KO and WT interneurons, aligned in the same way as in (F-H), respectively. *p < 0.05; **p < 0.01; ns, not significant. The statistical data are listed in Supplementary file 1– Supplementary Table 1. Data are mean ± SEM.

**Figure 7—figure supplement 1.**
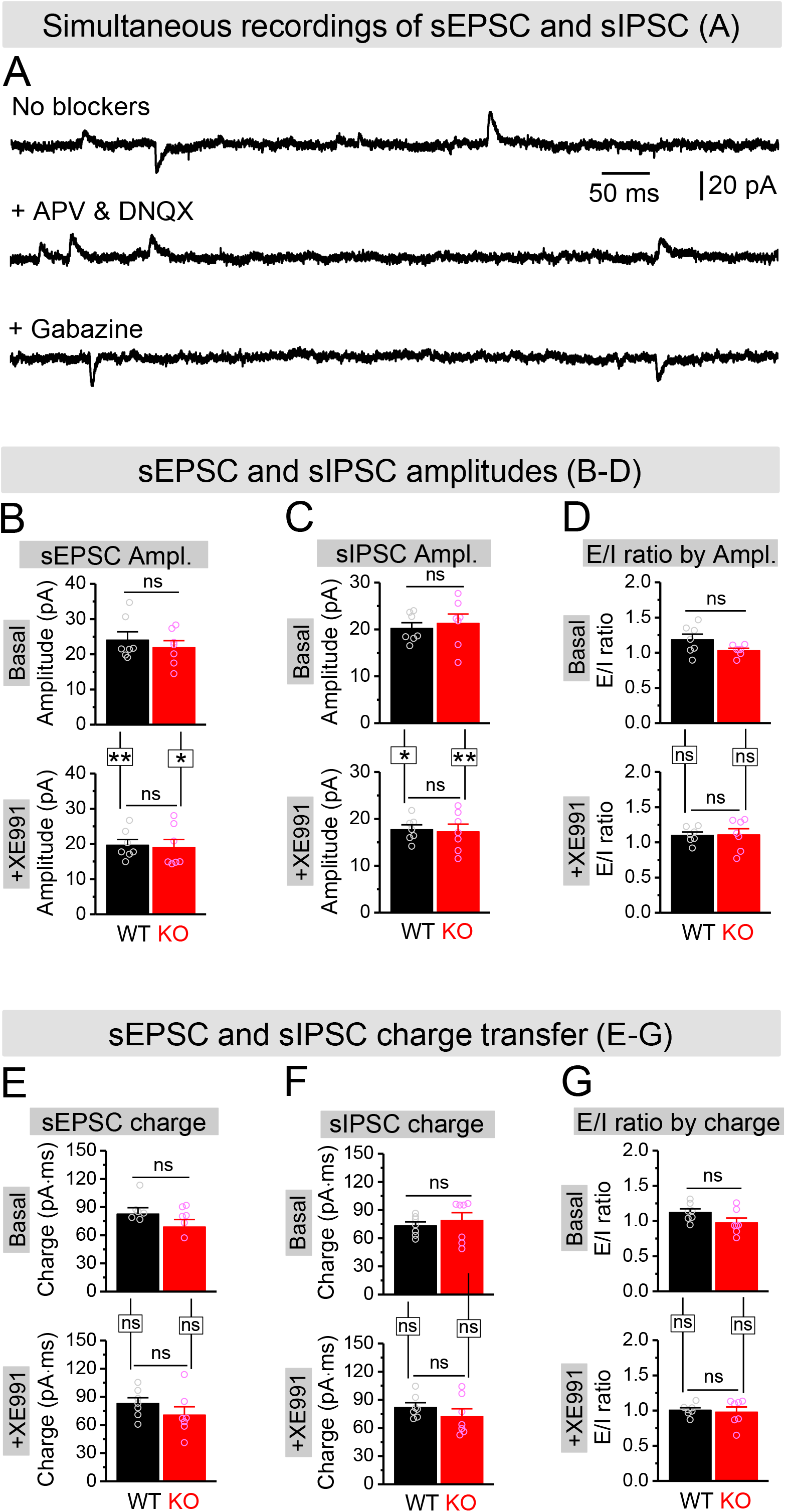
Effect of XE991 on spontaneous synaptic inputs onto granule cells. (A) Example traces for simultaneous recording of sEPSC and sIPSC from granule cells (holding at -40 mV). In the absence of blockers against glutamate and GABA ionotropic receptors, both sEPSC (down-ward) and sIPSC (up-ward) were exhibited (*upper trace*). In the presence of APV and DNQX (blockade of NMDA and AMPA receptors), only up-ward sIPSC was kept (*middle trace*). In contrast, in the presence of gabazine (blockade of GABA_A_ receptors), only down-ward sEPSC was observed (*lower trace*). (B and C) Amplitudes of simultaneously recorded sEPSC (B) and sIPSC (C) before (Basal, *upper bars*) and during XE991 (+ XE991, *lower bars*). (D) E/I ratio evaluated by amplitude before (Basal, *upper bars*) and during XE991 (+ XE991, *lower bars*). (E and F) The same as (B and C), but for charge transfers of simultaneously recorded sEPSC (E) and sIPSC (F). (G) The same as (D), but evaluated by charge transfer. *p < 0.05; **p < 0.01; ns, not significant. Horizontal lines denote comparison between genotypes; vertical lines indicate comparison between before and during XE991 within genotypes. The statistical data are listed in Supplementary file 1–Supplementary Table 1. Data are mean ± SEM.

**Figure 7—figure supplement 2.**
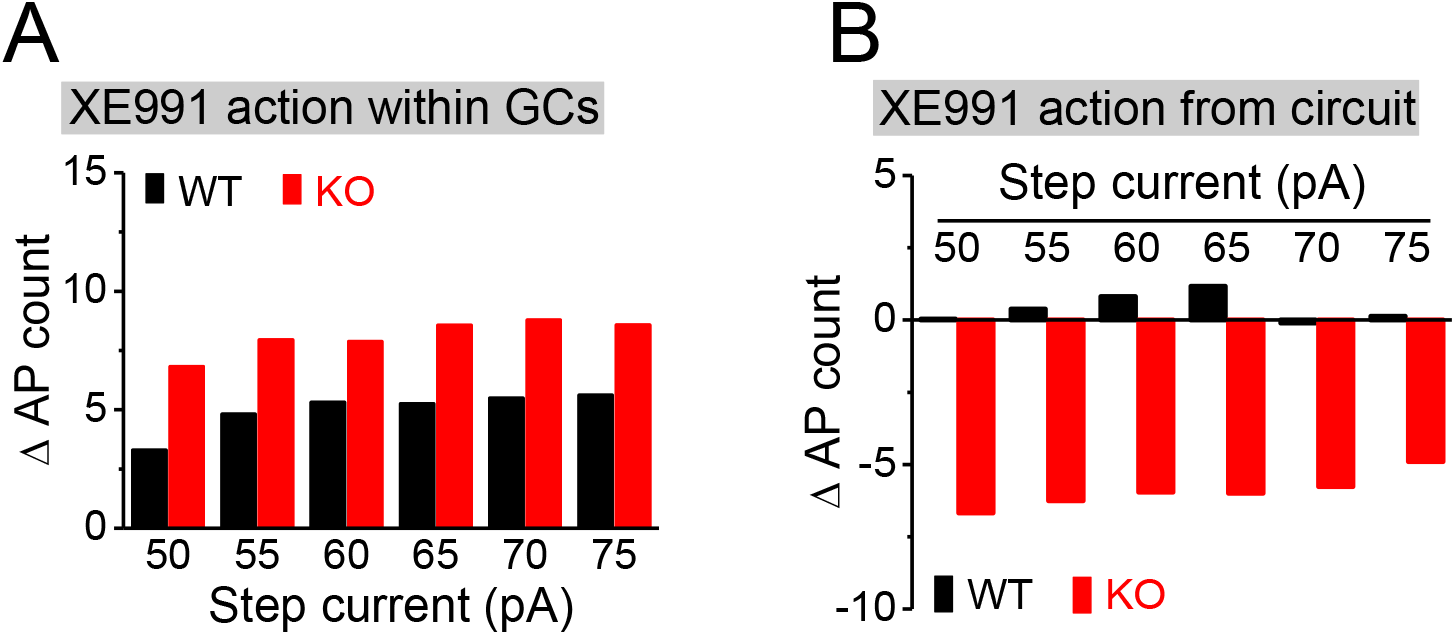
Estimation of direct and circuit effects of XE991 on granule cell excitability. (A) XE991 can directly increase excitability of granule cells, leading to increase AP firing. The direct effect of XE991 on AP firing were defined as the differences between before and during XE991 in the isolated granule cells (Figure 7F), ie, values of (7F*_lower_* – 7F*_upper_*), where 7F*_lower_*and 7F*_upper_* are the mean values from Figure 7F lower and upper panels, respectively. Note that XE991 directly increases number of AP largely independent of step current intensity in both genotypes, but with relative larger effect in KO mice. GCs, granule cells. (B) XE991 can also modulate granule cell excitability to change AP firing via its circuit effect. Values were estimated from the differences before and during XE991, between isolated granule cells (Figure 7F) and granule cells with intact circuit (Figure 7H), ie, values of [(7H*_lower_* – 7H*_upper_*) - (7F*_lower_* – 7F*_upper_*)], where 7F and 7H are the mean values from Figures 7F and 7H (lower or upper panel, accordingly). Note that the circuit effect of XE991 on AP firing was also independent of step current intensity (little effect in WT mice as shown by values fluctuating around 0; but dampening ∼6 APs in KO mice).

**Figure 8—figure supplement 1.**
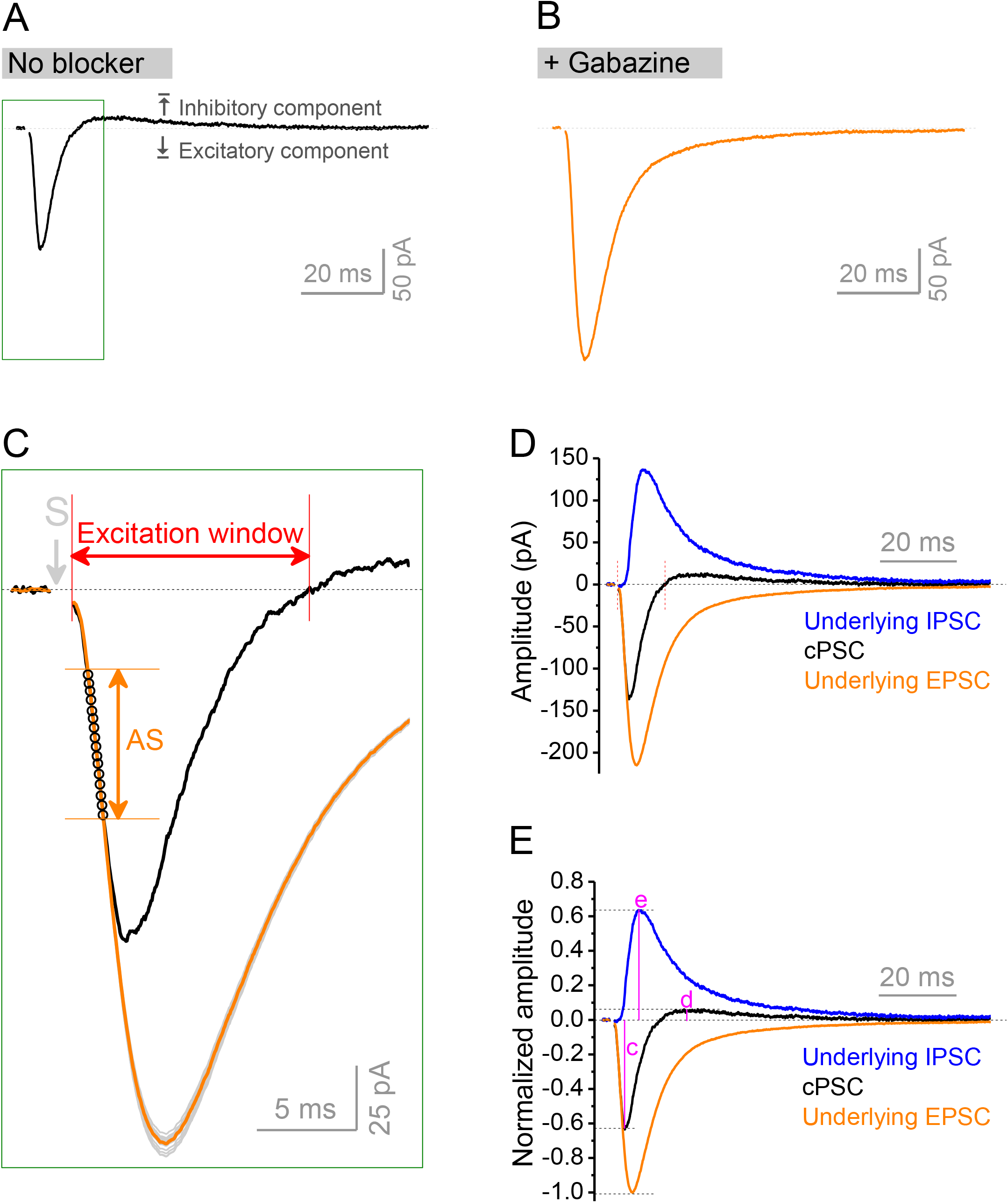
Isolation of underlying EPSC and IPSC from compound PSC. (A and B) Compound postsynaptic current (cPSC) was recorded from granule cells by holding membrane potential at -45 mV. The cPSC (A) shows an initial excitatory component (down-ward) and followed by an inhibitory component (up-ward). Dotted line denotes the baseline before stimulation. At the end of each recording, the pure EPSC (B) was recorded by keeping the same stimulation intensity and in the presence of gabazine, which was used to create EPSC template (an average of at least 20 uncontaminated EPSCs) for the same cell. Note that the amplitude and decay trajectory of pure EPSC was largely different from excitatory component of cPSC. Stimulation artifacts were removed. Boxed area in (A) was enlarged to show the procedure of approximation of underlying EPSC. (C) Zoom-in of boxed area in (A) showing approximation of underlying EPSC. The EPSC template was repeatedly scaled to each data point (black circles) of cPSC “approximating segment” (AS) to obtain a set of scaled EPSCs (gray traces). All scaled EPSCs were then averaged to approximate an underlying EPSC (orange trace) for a given cPSC (black trace). The cPSC “approximating segment” was defined as 25–65% height of cPSC excitatory component (but not beyond 2.5 ms after stimulation). Excitation window was defined as the full duration of cPSC excitatory component. Stimulation artifact was removed. “S” denotes the stimulation time point. (D) The underlying IPSC (blue trace) was isolated by subtracting the underlying EPSC (orange trace as that in C) from corresponding cPSC (black trace). For the purpose of view, the baseline before stimulation was shifted to be 0 in the figure. The two vertical red lines delimit the excitation window. (E) Normalized traces of cPSC, underlying EPSC and IPSC. For better comparison, we normalized the cPSC and underlying IPSC to their respective underlying EPSC, which reflects the PP stimulation intensity. The underlying EPSC and IPSC were then used to estimate E/I ratio by their peak amplitudes or charge transfers (amplitude-time integration within 100 ms). The letters **c**, **d** and **e** indicate the normalized peak amplitudes of cPSC excitatory, inhibitory components and underlying IPSC, which then summarized in Figures 8C, 8D and 8E, respectively.

**Figure 8—figure supplement 2.**
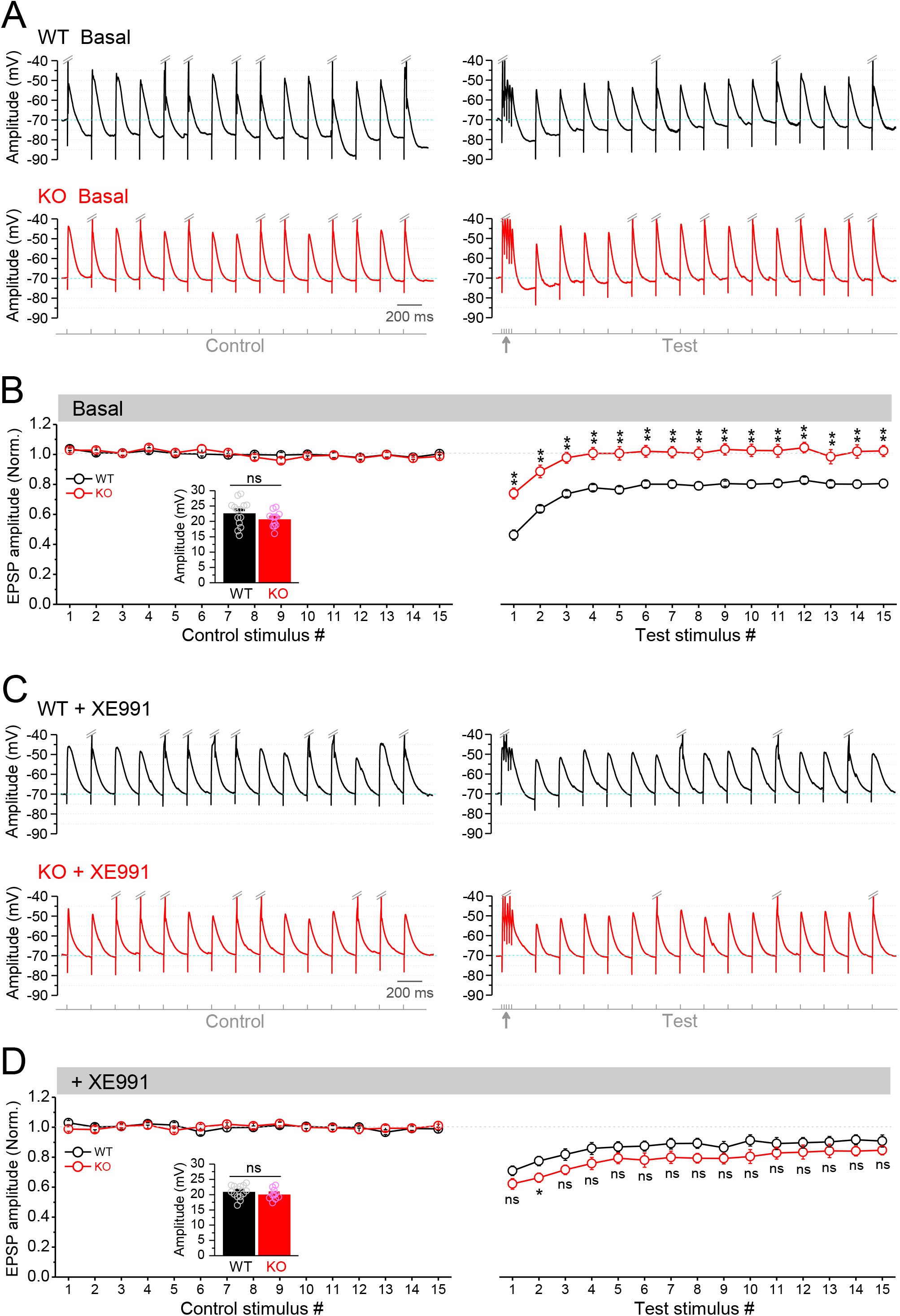
Circuit-wide inhibition of Kv7 channels enhanced the gamma burst-induced suppression of EPSP integration in *Fmr1* KO mice. (A) Sample Traces of EPSP in response to theta-gamma coupling stimulation of PP. Traces were zoomed in vertically (ie, amplitude dimension) and APs were truncated (indicated by double-slash) to emphasize EPSP size. EPSC amplitude was defined by the voltage difference between EPSC peak and -70 mV level (cyan line). *Lowermost panel* shows the stimulation paradigm. Up-pointing arrow indicates gamma-stimulation. (B) Summary data for experiments exemplified in (A) showing stable EPSP amplitude in control stimulation (*left panel*) and gamma-suppression of EPSP amplitude (*right panel*). *Insert,* real EPSP amplitude in control stimulation. (C) The same as in (A), but in the presence of XE991. (D) The same as in (B), but in the presence of XE991. Note that XE991 significantly dampened EPSP amplitude in KO mice (*right panel*) and the EPSP were largely comparable between genotypes. *p < 0.05; **p < 0.01; ns, not significant. The statistical data are listed in Supplementary file 1– Supplementary Table 1. Data are mean ± SEM.

**Figure 8—figure supplement 3.**
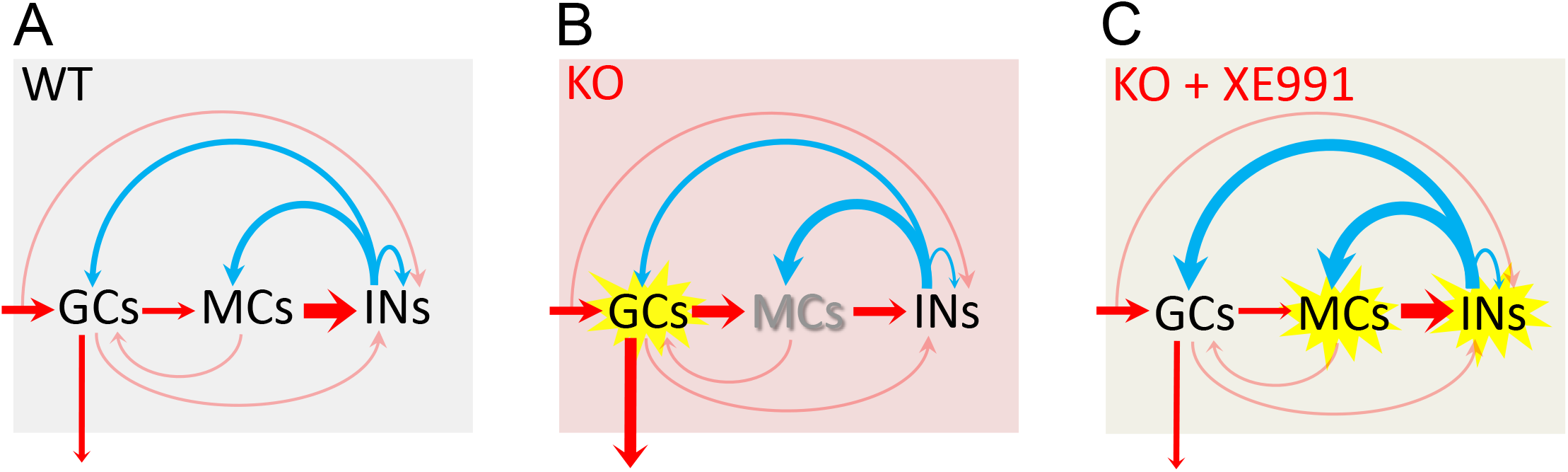
Circuit-wide inhibition of Kv7 channels corrected dentate gyrus output in *Fmr1* KO mice. (A) The diagram shows dentate gyrus output in response to theta-gamma coupling stimulation of PP in the WT mice. Here, we focus on the three-synapse indirect feedback inhibition pathway: Granule cells (GCs)➔MCs-+Interneurons (INs)-+GCs (thickness of arrows indicating synaptic drive weight). Because MCs (rather than granule cells) provide the main source of excitatory drive onto interneurons, the three-synapse indirect feedback inhibition likely surpasses the canonical two-synapse feedback inhibition (ie, GCs➔INs➔GCs). This model indicates that, in the normal condition, MCs integrate the incoming excitatory input and then relay and expand it to interneurons, which secure a sufficient and well-timed inhibition feedback onto granule cells to maintain a dynamically balanced E/I inputs onto granule cells and a narrow excitation window, and thus ensures sparse AP firing in granule cells. Red arrows designate excitation and blue arrows inhibition. The arrow entering shading areas from left side represents PP input (theta-gamma coupling stimulation). The downward arrow exiting the shading area indicates dentate output to CA3. The thinly semi-transparent arrows are synaptic connections whose contribution to cellular excitability has not been evaluated in the present study. The thickness of arrows denotes the synaptic drive weight of excitation (or inhibition). It is noteworthy that the actual synaptic drive weight of each synapse varies dynamically to maintain the precisely well-timed circuit E/I balance and then proper information processing. Thus, one may imagine that the thickness of each arrow (ie, synaptic drive weight) changes dynamically and sequentially (due to neurotransmission direction and synaptic delays) to understand dynamic E/I balance. (B) The same as in (**a**), but for KO mice. Loss of FMRP caused these changes (for details, see Figures 1, 4, 5, 6 and 8): 1) the granule cells are hyperexcitable (highlighted by yellow explosion marker), and dentate output are increased (thicker arrow compared with that of WT mice in A); 2) MCs are hypo-excitable (shadow font) and MCs output are decreased (thinner arrow); 3) both excitatory and inhibitory inputs onto MCs are increased (thicker arrows); and 4) both excitatory and inhibitory inputs onto INs are reduced (thinner arrows). (C) The same as in (B), but in the presence of XE991 (circuit-wide inhibition of Kv7 channels). XE991 caused these changes in the KO mice (for details, see Figure 8 and Figure 8—figure supplement 2): 1) Owing to the abnormal enhanced Kv7 function in KO MCs, XE991 boosted up MC excitability (explosion marker) to enhance excitatory output (thicker red arrow) onto interneurons, which increases interneurons excitability (explosion marker); and 2) XE991 increased inhibitory drive onto granule cells (thicker blue arrow onto granule cells compared with that of in (B), which dampens EPSP integration (due to summation of enhanced inhibitory input) in granule cells and suppresses dentate gyrus output.

**Supplementary File 1**. Supplementary Table 1. Table containing statistical information for all figures.

## Notes

### Competing Interest Statement

The authors have declared no competing interest.

### Summary of Updates

This version of the manuscript has been revised to address the reviewers' concerns to update the following: Results, Discussion, Methods, Figures, Figure Legends

## References

Amaral DG, Scharfman HE, Lavenex P. 2007. The dentate gyrus: fundamental neuroanatomical organization (dentate gyrus for dummies). Prog Brain Res 163: 3–22

Antoine MW, Langberg T, Schnepel P, Feldman DE. 2019. Increased Excitation-Inhibition Ratio Stabilizes Synapse and Circuit Excitability in Four Autism Mouse Models. Neuron 101: 648–61 e4

Baculis BC, Zhang J, Chung HJ. 2020. The Role of Kv7 Channels in Neural Plasticity and Behavior. Front Physiol 11: 568667

Bartley AF, Dobrunz LE. 2015. Short-term plasticity regulates the excitation/inhibition ratio and the temporal window for spike integration in CA1 pyramidal cells. Eur J Neurosci 41: 1402–15

Berron D, Schutze H, Maass A, Cardenas-Blanco A, Kuijf HJ, Kumaran D, Duzel E. 2016. Strong Evidence for Pattern Separation in Human Dentate Gyrus. J Neurosci 36: 7569–79

Bhatia A, Moza S, Bhalla US. 2021. Precise excitation-inhibition balance controls gain and timing in the hippocampus. Elife 8

Booker SA, Domanski APF, Dando OR, Jackson AD, Isaac JTR, et al. 2019. Altered dendritic spine function and integration in a mouse model of fragile X syndrome. Nat Commun 10: 4813

Bostrom C, Yau SY, Majaess N, Vetrici M, Gil-Mohapel J, Christie BR. 2016. Hippocampal dysfunction and cognitive impairment in Fragile-X Syndrome. Neurosci Biobehav Rev 68: 563–74

Bott JB, Muller MA, Jackson J, Aubert J, Cassel JC, Mathis C, Goutagny R. 2016. Spatial Reference Memory is Associated with Modulation of Theta-Gamma Coupling in the Dentate Gyrus. Cereb Cortex 26: 3744–53

Botterill JJ, Gerencer KJ, Vinod KY, Alcantara-Gonzalez D, Scharfman HE. 2021a. Dorsal and ventral mossy cells differ in their axonal projections throughout the dentate gyrus of the mouse hippocampus. Hippocampus 31: 522–39

Botterill JJ, Vinod KY, Gerencer KJ, Teixeira CM, LaFrancois JJ, Scharfman HE. 2021b. Bidirectional Regulation of Cognitive and Anxiety-like Behaviors by Dentate Gyrus Mossy Cells in Male and Female Mice. J Neurosci 41: 2475–95

Brown DA, Passmore GM. 2009. Neural KCNQ (Kv7) channels. Br J Pharmacol 156: 1185–95

Bui AD, Nguyen TM, Limouse C, Kim HK, Szabo GG, et al. 2018. Dentate gyrus mossy cells control spontaneous convulsive seizures and spatial memory. Science 359: 787–90

Butler CR, Westbrook GL, Schnell E. 2022. Adaptive Mossy Cell Circuit Plasticity after Status Epilepticus. J Neurosci 42: 3025–36

Chancey JH, Poulsen DJ, Wadiche JI, Overstreet-Wadiche L. 2014. Hilar mossy cells provide the first glutamatergic synapses to adult-born dentate granule cells. J Neurosci 34: 2349–54

Chiu CQ, Castillo PE. 2008. Input-specific plasticity at excitatory synapses mediated by endocannabinoids in the dentate gyrus. Neuropharmacology 54: 68–78

Contractor A, Klyachko VA, Portera-Cailliau C. 2015. Altered Neuronal and Circuit Excitability in Fragile X Syndrome. Neuron 87: 699–715

Csicsvari J, Jamieson B, Wise KD, Buzsaki G. 2003. Mechanisms of gamma oscillations in the hippocampus of the behaving rat. Neuron 37: 311–22

Darnell JC, Van Driesche SJ, Zhang C, Hung KY, Mele A, et al. 2011. FMRP stalls ribosomal translocation on mRNAs linked to synaptic function and autism. Cell 146: 247–61

Delmas P, Brown DA. 2005. Pathways modulating neural KCNQ/M (Kv7) potassium channels. Nat Rev Neurosci 6: 850–62

Deng PY, Avraham O, Cavalli V, Klyachko VA. 2021. Hyperexcitability of Sensory Neurons in Fragile X Mouse Model. Front Mol Neurosci 14: 796053

Deng PY, Carlin D, Oh YM, Myrick LK, Warren ST, Cavalli V, Klyachko VA. 2019. Voltage-Independent SK-Channel Dysfunction Causes Neuronal Hyperexcitability in the Hippocampus of Fmr1 Knock-Out Mice. J Neurosci 39: 28–43

Deng PY, Klyachko VA. 2016. Increased Persistent Sodium Current Causes Neuronal Hyperexcitability in the Entorhinal Cortex of Fmr1 Knockout Mice. Cell Rep 16: 3157–66

Deng PY, Klyachko VA. 2021. Channelopathies in fragile X syndrome. Nat Rev Neurosci 22: 275–89

Deng PY, Kumar A, Cavalli V, Klyachko VA. 2022. FMRP regulates GABAA receptor channel activity to control signal integration in hippocampal granule cells. Cell Rep 39: 110820

Deng PY, Rotman Z, Blundon JA, Cho Y, Cui J, et al. 2013. FMRP regulates neurotransmitter release and synaptic information transmission by modulating action potential duration via BK channels. Neuron 77: 696–711

Doll CA, Broadie K. 2014. Impaired activity-dependent neural circuit assembly and refinement in autism spectrum disorder genetic models. Front Cell Neurosci 8: 30

Domanski APF, Booker SA, Wyllie DJA, Isaac JTR, Kind PC. 2019. Cellular and synaptic phenotypes lead to disrupted information processing in Fmr1-KO mouse layer 4 barrel cortex. Nat Commun 10: 4814

Eadie BD, Cushman J, Kannangara TS, Fanselow MS, Christie BR. 2012. NMDA receptor hypofunction in the dentate gyrus and impaired context discrimination in adult Fmr1 knockout mice. Hippocampus 22: 241–54

Fredes F, Shigemoto R. 2021. The role of hippocampal mossy cells in novelty detection. Neurobiol Learn Mem 183: 107486

Ghilan M, Hryciw BN, Brocardo PS, Bostrom CA, Gil-Mohapel J, Christie BR. 2015. Enhanced corticosteroid signaling alters synaptic plasticity in the dentate gyrus in mice lacking the fragile X mental retardation protein. Neurobiol Dis 77: 26–34

Gibson JR, Bartley AF, Hays SA, Huber KM. 2008. Imbalance of neocortical excitation and inhibition and altered UP states reflect network hyperexcitability in the mouse model of fragile X syndrome. J Neurophysiol 100: 2615–26

Gilling M, Rasmussen HB, Calloe K, Sequeira AF, Baretto M, et al. 2013. Dysfunction of the Heteromeric KV7.3/KV7.5 Potassium Channel is Associated with Autism Spectrum Disorders. Front Genet 4: 54

Golomb D, Donner K, Shacham L, Shlosberg D, Amitai Y, Hansel D. 2007. Mechanisms of firing patterns in fast-spiking cortical interneurons. PLoS Comput Biol 3: e156

Grangeray-Vilmint A, Valera AM, Kumar A, Isope P. 2018. Short-Term Plasticity Combines with Excitation-Inhibition Balance to Expand Cerebellar Purkinje Cell Dynamic Range. J Neurosci 38: 5153–67

Greene DL, Hoshi N. 2017. Modulation of Kv7 channels and excitability in the brain. Cell Mol Life Sci 74: 495–508

Gu N, Vervaeke K, Hu H, Storm JF. 2005. Kv7/KCNQ/M and HCN/h, but not KCa2/SK channels, contribute to the somatic medium after-hyperpolarization and excitability control in CA1 hippocampal pyramidal cells. J Physiol 566: 689–715

Haick JM, Brueggemann LI, Cribbs LL, Denning MF, Schwartz J, Byron KL. 2017. PKC-dependent regulation of Kv7.5 channels by the bronchoconstrictor histamine in human airway smooth muscle cells. Am J Physiol Lung Cell Mol Physiol 312: L822–L34

Hashimotodani Y, Nasrallah K, Jensen KR, Chavez AE, Carrera D, Castillo PE. 2017. LTP at Hilar Mossy Cell-Dentate Granule Cell Synapses Modulates Dentate Gyrus Output by Increasing Excitation/Inhibition Balance. Neuron 95: 928–43 e3

Hasselmo ME, Giocomo LM, Zilli EA. 2007. Grid cell firing may arise from interference of theta frequency membrane potential oscillations in single neurons. Hippocampus 17: 1252–71

Houser CR, Peng Z, Wei X, Huang CS, Mody I. 2021. Mossy Cells in the Dorsal and Ventral Dentate Gyrus Differ in Their Patterns of Axonal Projections. J Neurosci 41: 991–1004

Incontro S, Sammari M, Azzaz F, Inglebert Y, Ankri N, et al. 2021. Endocannabinoids Tune Intrinsic Excitability in O-LM Interneurons by Direct Modulation of Postsynaptic Kv7 Channels. J Neurosci 41: 9521–38

Jentsch TJ. 2000. Neuronal KCNQ potassium channels: physiology and role in disease. Nat Rev Neurosci 1: 21–30

Jinde S, Zsiros V, Jiang Z, Nakao K, Pickel J, et al. 2012. Hilar mossy cell degeneration causes transient dentate granule cell hyperexcitability and impaired pattern separation. Neuron 76: 1189–200

Jinde S, Zsiros V, Nakazawa K. 2013. Hilar mossy cell circuitry controlling dentate granule cell excitability. Front Neural Circuits 7: 14

Jones F, Gamper N, Gao H. 2021. Kv7 Channels and Excitability Disorders. Handb Exp Pharmacol 267: 185–230

Kennedy T, Rinker D, Broadie K. 2020. Genetic background mutations drive neural circuit hyperconnectivity in a fragile X syndrome model. BMC Biol 18: 94

Lee B, Panda S, Lee HY. 2020. Primary Ciliary Deficits in the Dentate Gyrus of Fragile X Syndrome. Stem Cell Reports 15: 454–66

Leutgeb JK, Leutgeb S, Moser MB, Moser EI. 2007. Pattern separation in the dentate gyrus and CA3 of the hippocampus. Science 315: 961–6

Lisman JE, Jensen O. 2013. The theta-gamma neural code. Neuron 77: 1002–16

Liu Y, Bian X, Wang K. 2021. Pharmacological Activation of Neuronal Voltage-Gated Kv7/KCNQ/M-Channels for Potential Therapy of Epilepsy and Pain. Handb Exp Pharmacol 267: 231–51

Lubke J, Frotscher M, Spruston N. 1998. Specialized electrophysiological properties of anatomically identified neurons in the hilar region of the rat fascia dentata. J Neurophysiol 79: 1518–34

Marsillo A, David L, Gerges B, Kerr D, Sadek R, et al. 2021. PKC epsilon as a neonatal target to correct FXS-linked AMPA receptor translocation in the hippocampus, boost PVN oxytocin expression, and normalize adult behavior in Fmr1 knockout mice. Biochim Biophys Acta Mol Basis Dis 1867: 166048

Martinello K, Huang Z, Lujan R, Tran B, Watanabe M, et al. 2015. Cholinergic afferent stimulation induces axonal function plasticity in adult hippocampal granule cells. Neuron 85: 346–63

Miceli F, Soldovieri MV, Ambrosino P, De Maria M, Migliore M, Migliore R, Taglialatela M. 2015. Early-onset epileptic encephalopathy caused by gain-of-function mutations in the voltage sensor of Kv7.2 and Kv7.3 potassium channel subunits. J Neurosci 35: 3782–93

Miceli F, Soldovieri MV, Martire M, Taglialatela M. 2008. Molecular pharmacology and therapeutic potential of neuronal Kv7-modulating drugs. Curr Opin Pharmacol 8: 65–74

Mircheva Y, Peralta MR, 3rd, Toth K. 2019. Interplay of Entorhinal Input and Local Inhibitory Network in the Hippocampus at the Origin of Slow Inhibition in Granule Cells. J Neurosci 39: 6399–413

Mizuseki K, Sirota A, Pastalkova E, Buzsaki G. 2009. Theta oscillations provide temporal windows for local circuit computation in the entorhinal-hippocampal loop. Neuron 64: 267–80

Modgil A, Vien TN, Ackley MA, Doherty JJ, Moss SJ, Davies PA. 2019. Neuroactive Steroids Reverse Tonic Inhibitory Deficits in Fragile X Syndrome Mouse Model. Front Mol Neurosci 12: 15

Monday HR, Kharod SC, Yoon YJ, Singer RH, Castillo PE. 2022. Presynaptic FMRP and local protein synthesis support structural and functional plasticity of glutamatergic axon terminals. Neuron 110: 2588–606 e6

Mott DD, Turner DA, Okazaki MM, Lewis DV. 1997. Interneurons of the dentate-hilus border of the rat dentate gyrus: morphological and electrophysiological heterogeneity. J Neurosci 17: 3990–4005

Nappi P, Miceli F, Soldovieri MV, Ambrosino P, Barrese V, Taglialatela M. 2020. Epileptic channelopathies caused by neuronal Kv7 (KCNQ) channel dysfunction. Pflugers Arch 472: 881–98

Neves L, Lobao-Soares B, Araujo APC, Furtunato AMB, Paiva I, et al. 2022. Theta and gamma oscillations in the rat hippocampus support the discrimination of object displacement in a recognition memory task. Front Behav Neurosci 16: 970083

Patel AB, Hays SA, Bureau I, Huber KM, Gibson JR. 2013. A target cell-specific role for presynaptic Fmr1 in regulating glutamate release onto neocortical fast-spiking inhibitory neurons. J Neurosci 33: 2593–604

Patel AB, Loerwald KW, Huber KM, Gibson JR. 2014. Postsynaptic FMRP promotes the pruning of cell-to-cell connections among pyramidal neurons in the L5A neocortical network. J Neurosci 34: 3413–8

Pelkey KA, Chittajallu R, Craig MT, Tricoire L, Wester JC, McBain CJ. 2017. Hippocampal GABAergic Inhibitory Interneurons. Physiol Rev 97: 1619–747

Pernia-Andrade AJ, Jonas P. 2014. Theta-gamma-modulated synaptic currents in hippocampal granule cells in vivo define a mechanism for network oscillations. Neuron 81: 140–52

Peters HC, Hu H, Pongs O, Storm JF, Isbrandt D. 2005. Conditional transgenic suppression of M channels in mouse brain reveals functions in neuronal excitability, resonance and behavior. Nat Neurosci 8: 51–60

Remmers CL, Contractor A. 2018. Development of GABAergic Inputs Is Not Altered in Early Maturation of Adult Born Dentate Granule Neurons in Fragile X Mice. eNeuro 5

Salcedo-Arellano MJ, Hagerman RJ, Martinez-Cerdeno V. 2020. Fragile X syndrome: clinical presentation, pathology and treatment. Gac Med Mex 156: 60–66

Sathyanarayana SH, Saunders JA, Slaughter J, Tariq K, Chakrabarti R, et al. 2022. Pten heterozygosity restores neuronal morphology in fragile X syndrome mice. Proc Natl Acad Sci U S A 119: e2109448119

Scharfman HE. 2016. The enigmatic mossy cell of the dentate gyrus. Nat Rev Neurosci 17: 562–75

Scharfman HE. 2018. Advances in understanding hilar mossy cells of the dentate gyrus. Cell Tissue Res 373: 643–52

Scharfman HE, Myers CE. 2013. Hilar mossy cells of the dentate gyrus: a historical perspective. Front Neural Circuits 6: 106

Sears JC, Broadie K. 2018. Fragile X Mental Retardation Protein Regulates Activity-Dependent Membrane Trafficking and Trans-Synaptic Signaling Mediating Synaptic Remodeling. Front Mol Neurosci 10: 440

Shigemoto R, Kinoshita A, Wada E, Nomura S, Ohishi H, et al. 1997. Differential presynaptic localization of metabotropic glutamate receptor subtypes in the rat hippocampus. J Neurosci 17: 7503–22

Sloviter RS, Bumanglag AV, Schwarcz R, Frotscher M. 2012. Abnormal dentate gyrus network circuitry in temporal lobe epilepsy.

Springer K, Varghese N, Tzingounis AV. 2021. Flexible Stoichiometry: Implications for KCNQ2-and KCNQ3-Associated Neurodevelopmental Disorders. Dev Neurosci 43: 191–200

Tessier CR, Broadie K. 2009. Activity-dependent modulation of neural circuit synaptic connectivity. Front Mol Neurosci 2: 8

Tzingounis AV, Heidenreich M, Kharkovets T, Spitzmaul G, Jensen HS, Nicoll RA, Jentsch TJ. 2010. The KCNQ5 potassium channel mediates a component of the afterhyperpolarization current in mouse hippocampus. Proc Natl Acad Sci U S A 107: 10232–7

van der Horst J, Greenwood IA, Jepps TA. 2020. Cyclic AMP-Dependent Regulation of Kv7 Voltage-Gated Potassium Channels. Front Physiol 11: 727

Wang KY, Wu JW, Cheng JK, Chen CC, Wong WY, et al. 2021. Elevation of hilar mossy cell activity suppresses hippocampal excitability and avoidance behavior. Cell Rep 36: 109702

Willemsen R, Kooy RF. 2017. Fragile X Syndrome: From Genetics to Targeted Treatment. Elsevier. 498 pp.

Yau SY, Bettio L, Chiu J, Chiu C, Christie BR. 2019. Fragile-X Syndrome Is Associated With NMDA Receptor Hypofunction and Reduced Dendritic Complexity in Mature Dentate Granule Cells. Front Mol Neurosci 11: 495

Yau SY, Bostrom CA, Chiu J, Fontaine CJ, Sawchuk S, et al. 2016. Impaired bidirectional NMDA receptor dependent synaptic plasticity in the dentate gyrus of adult female Fmr1 heterozygous knockout mice. Neurobiol Dis 96: 261–70

Yeh CY, Asrican B, Moss J, Quintanilla LJ, He T, et al. 2018. Mossy Cells Control Adult Neural Stem Cell Quiescence and Maintenance through a Dynamic Balance between Direct and Indirect Pathways. Neuron 99: 493–510 e4

Yun SH, Trommer BL. 2011. Fragile X mice: reduced long-term potentiation and N-Methyl-D-Aspartate receptor-mediated neurotransmission in dentate gyrus. J Neurosci Res 89: 176–82

Zhou S, Yu Y. 2018. Synaptic E-I Balance Underlies Efficient Neural Coding. Front Neurosci 12: 46

