## Supplemental Table 1 for "Circuit-based intervention corrects excessive dentate gyrus output in the Fragile X mouse model"

**Supplementary file 1—Supplementary Table 1.** Statistical data for figures

**Figure 1.** Decreased excitability of dentate mossy cells in *Fmr1* KO mice

p < 0.05 in red

| Figure 1 | Condition | Genotype | mean ± SEM | Number of cells tested | p value (WT vs KO) |
|---|---|---|---|---|---|
| **Figure 1C** AP count (/20 s) | -64 mV | WT | 0.75 ± 0.25 | 8 | 0.16886 |
| | | KO | 0.29 ± 0.18 | 7 | |
| | -63 mV | WT | 1.00 ± 0.38 | 8 | 0.59620 |
| | | KO | 0.71 ± 0.36 | 7 | |
| | -62 mV | WT | 1.38 ± 0.50 | 8 | 0.35773 |
| | | KO | 0.71 ± 0.47 | 7 | |
| | -61 mV | WT | 3.00 ± 0.65 | 8 | **0.01015** |
| | | KO | 0.57 ± 0.43 | 7 | |
| | -60 mV | WT | 4.00 ± 1.21 | 8 | **0.04443** |
| | | KO | 0.86 ± 0.59 | 7 | |
| | -59 mV | WT | 8.63 ± 2.12 | 8 | **0.00783** |
| | | KO | 1.29 ± 0.57 | 7 | |
| | -58 mV | WT | 12.38 ± 2.59 | 8 | **0.00331** |
| | | KO | 2.00 ± 0.85 | 7 | |
| | -57 mV | WT | 17.88 ± 2.21 | 8 | **0.00018** |
| | | KO | 3.71 ± 1.48 | 7 | |
| | -56 mV | WT | 23.88 ± 3.48 | 8 | **0.02750** |
| | | KO | 9.14 ± 4.95 | 7 | |
| | -55 mV | WT | 30.75 ± 4.71 | 8 | **0.04470** |
| | | KO | 14.71 ± 5.53 | 7 | |

| Figure 1 | Parameter | Genotype | mean ± SEM | Number of cells tested | p value (WT vs KO) |
|---|---|---|---|---|---|
| **Figure 1E** AP | Count (/2-s ramp) | WT | 16.80 ± 2.66 | 18 | **0.01970** |
| | | KO | 9.97 ± 1.02 | 19 | |
| **Figure 1F** AP | Threshold (mV) | WT | -45.06 ± 0.78 | 18 | **0.00318** |
| | | KO | -42.14 ± 0.51 | 19 | |
| **Figure 1G** AP | Rheobase (pA) | WT | 152.58 ± 11.66 | 18 | **0.00020** |
| | | KO | 211.42 ± 7.82 | 19 | |
| **Figure 1H** AP | Rheo-charge (pC) | WT | 81.77 ± 12.00 | 18 | **0.00020** |
| | | KO | 150.44 ± 10.92 | 19 | |
| **Figure 1I** AP | Max. rise rate (mV/ms) | WT | 300.73 ± 11.12 | 16 | 0.26123 |
| | | KO | 283.67 ± 9.16 | 13 | |
| **Figure 1J** AP | Rise time (ms) | WT | 0.267 ± 0.007 | 16 | 0.78010 |
| | | KO | 0.269 ± 0.005 | 13 | |
| **Figure 1K** AP | Fall time (ms) | WT | 0.677 ± 0.023 | 16 | 0.76199 |
| | | KO | 0.687 ± 0.018 | 13 | |
| **Figure 1L** AP | Duration (ms) | WT | 0.714 ± 0.022 | 16 | 0.56462 |
| | | KO | 0.73 ± 0.013 | 13 | |
| **Figure 1M** AP | Peak potential (mV) | WT | 34.63 ± 1.98 | 16 | 0.90338 |
| | | KO | 34.94 ± 1.39 | 13 | |
| **Figure 1N** AP | Ampliutde (mV) | WT | 79.88 ± 1.24 | 16 | 0.09566 |
| | | KO | 76.72 ± 1.34 | 13 | |

**Figure 2.** Decreased input resistance around threshold potential in Fmr1 KO mossy cells

| Figure 2 | Parameter | Genotype | mean ± SEM | Number of cells tested | p value (WT vs KO) | p value Before XE vs during XE within genotype |
|---|---|---|---|---|---|---|
| **Figure 2A** | RMP (mV) | WT | -63.38 ± 1.15 | 13 | 0.34397 | N/A |
| | | KO | -65.09 ± 1.35 | 13 | | |
| **Figure 2B** | Capacitance (pF) | WT | 132.00 ± 12.00 | 11 | 0.73345 | N/A |
| | | KO | 137.73 ± 11.16 | 11 | | |
| **Figure 2C** @ RMP | Input resistance (MΩ) | WT | 167.13 ± 19.49 | 8 | 0.91027 | N/A |
| | | KO | 164.23 ± 15.13 | 9 | | |
| **Figure 2D** @ -45 mV (No XE) | Input resistance (MΩ) | WT | 90.23 ± 4.67 | 7 | **0.03261** | N/A |
| | | KO | 75.93 ± 3.82 | 8 | | |

| | | | | | | | |
|---|---|---|---|---|---|---|---|
| **Figure 2E** @ -45 mV +XE991 | Input resistance (MΩ) | WT | 96.67 ± 4.83 | 7 | 0.39087 | WT (No XE vs + XE) | **0.00034** |
| | | KO | 91.18 ± 3.96 | 8 | | KO (No XE vs + XE) | **0.00068** |
| **Figure 2F** | Input resistance change (MΩ) | WT | 6.44 ± 0.88 | 7 | **0.01062** | N/A | |
| | | KO | 15.25 ± 2.64 | 8 | | | |

**Figure 3.** Enhanced Kv7 function causes hypo-excitability in Fmr1 KO mossy cells

| Figure 3 | Parameter | Genotype | mean ± SEM | Number of cells tested | p value (WT vs KO) |
|---|---|---|---|---|---|
| **Figure 3A** | Δ holding current (pA) | WT | -20.13 ± 4.57 | 7 | **0.04644** |
| | | KO | -39.44 ± 7.79 | 4 | |
| **Figure 3B** | Δ membrane potential (mV) | WT | 0.38 ± 0.32 | 7 | **0.00011** |
| | | KO | 2.24 ± 0.20 | 10 | |
| **Figure 3D** | Kv7 current (pA/pF) | WT | 0.16 ± 0.04 | 6 | **0.02328** |
| | | KO | 0.32 ± 0.04 | 5 | |

| Figure 3 | Condition | Genotype | mean ± SEM | Number of cells tested | p value (WT vs KO) |
|---|---|---|---|---|---|
| **Figure 3E** AP Threshold (mV) | Isolated MCs | WT | -43.73 ± 0.47 | 7 | **0.00616** |
| | | KO | -41.50 ± 0.49 | 8 | |
| | Isolated MCs + XE991 | WT | -43.91 ± 0.31 | 7 | 0.51203 |
| | | KO | -43.48 ± 0.53 | 8 | |
| **Figure 3F** AP Rheobase (pA) | Isolated MCs | WT | 157.98 ± 9.18 | 7 | **0.00937** |
| | | KO | 204.79 ± 11.89 | 8 | |
| | Isolated MCs + XE991 | WT | 151.22 ± 7.31 | 7 | 0.32481 |
| | | KO | 164.45 ± 10.24 | 8 | |
| **Figure 3G** Rheobase charge transfer (pC) | Isolated MCs | WT | 86.81 ± 6.42 | 7 | **0.00646** |
| | | KO | 143.18 ± 15.22 | 8 | |
| | Isolated MCs + XE991 | WT | 76.14 ± 6.62 | 7 | 0.22376 |
| | | KO | 92.77 ± 10.68 | 8 | |
| **Figure 3H** AP count (/2-s ramp) | Isolated MCs | WT | 30.80 ± 4.87 | 7 | **0.01664** |
| | | KO | 16.48 ± 2.41 | 8 | |
| | Isolated MCs + XE991 | WT | 35.76 ± 4.08 | 7 | 0.11902 |
| | | KO | 26.68 ± 3.63 | 8 | |

**Figure 4.** Increased E/I ratio of inputs onto *Fmr1* KO mossy cells

| Figure 4 | Parameter | Genotype | mean ± SEM | Number of cells tested | p value (WT vs KO) |
|---|---|---|---|---|---|
| **Figure 4B** Mossy cells | Cumulative probability***** | WT | N/A | 7 | **0.00073** |
| | | KO | N/A | 8 | |
| | sEPSC count (/min) | WT | 795.86 ± 158.25 | 7 | **0.01058** |
| | | KO | 1504.13 ± 173.43 | 8 | |
| **Figure 4D** Mossy cells | Cumulative probability***** | WT | N/A | 18 | **0.03440** |
| | | KO | N/A | 22 | |
| | sIPSC count (/min) | WT | 527.33 ± 59.52 | 18 | **0.00207** |
| | | KO | 839.91 ± 70.22 | 22 | |
| **Figure 4E** Mossy cells | sEPSC amplitude (pA) | WT | 73.58 ± 9.33 | 7 | 0.27954 |
| | | KO | 90.11 ± 10.71 | 9 | |
| **Figure 4F** Mossy cells | sIPSC amplitude (pA) | WT | 45.74 ± 3.05 | 18 | 0.55236 |
| | | KO | 42.94 ± 3.40 | 22 | |
| **Figure 4G** Mossy cells | sEPSC Mean frequency (Hz) | WT | 13.26 ± 2.64 | 7 | **0.01058** |
| | | KO | 25.07 ± 2.89 | 8 | |
| **Figure 4H** Mossy cells | sIPSC Mean frequency (Hz) | WT | 8.79 ± 0.99 | 18 | **0.00207** |
| | | KO | 14.00 ± 1.17 | 22 | |
| **Figure 4I** Mossy cells | E/I ratio | WT | 1.51 | Estimated by mean values | |
| | | KO | 1.79 (↑18.66%) | | |

*, K-S test was used for comparison of cumulative probability. ↑ in KO E/I ratio indicating the percentage of the KO value increases or decreases relative to respective WT value.

**Figure 5.** Decreased E/I ratio of inputs onto *Fmr1* KO interneurons

| Figure 5 | Parameter | Genotype | mean ± SEM | Number of cells tested | p value (WT vs KO) |
|---|---|---|---|---|---|
| **Figure 5B** Interneurons | Cumulative probability* | WT | N/A | 8 | **<0.00001** |
| | | KO | N/A | 9 | |
| | sEPSC count (/min) | WT | 675.88 ± 128.88 | 8 | **0.00048** |
| | | KO | 119.00 ± 31.00 | 9 | |
| **Figure 5D** Interneurons | Cumulative probability* | WT | N/A | 11 | **<0.00001** |
| | | KO | N/A | 6 | |
| | sIPSC count (/min) | WT | 145.36 ± 21.83 | 11 | **0.01753** |
| | | KO | 61.33 ± 13.3 | 6 | |
| **Figure 5E** Interneurons | sEPSC amplitude (pA) | WT | 52.50 ± 4.68 | 8 | 0.38597 |
| | | KO | 43.75 ± 8.22 | 9 | |
| **Figure 5F** Interneurons | sIPSC amplitude (pA) | WT | 42.36 ± 5.24 | 11 | 0.25395 |
| | | KO | 33.11 ± 4.18 | 6 | |
| **Figure 5G** Interneurons | sEPSC Mean frequency (Hz) | WT | 11.26 ± 2.15 | 8 | **0.00048** |
| | | KO | 1.98 ± 0.52 | 9 | |
| **Figure 5H** Interneurons | sIPSC Mean frequency (Hz) | WT | 2.42 ± 0.36 | 11 | **0.01753** |
| | | KO | 1.02 ± 0.22 | 6 | |
| **Figure 5I** Interneurons | E/I ratio | WT | 4.65 | Estimated by mean values | |
| | | KO | 1.94 (↓58.27%) | | |

*, K-S test was used for comparison of cumulative probability. N/A, not applicable. ↓ in KO E/I ratio indicating the percentage of the KO value increases or decreases relative to respective WT value.

**Figure 6.** Mossy cells provide the main excitatory drive onto hilar interneurons

| Figure 6 | Genotype | Condition | mean ± SEM | Number of cells tested | p value (within genotype) [Basal vs +DCG-IV (or +WIN)] | |
|---|---|---|---|---|---|---|
| **Figure 6C** **Interneurons** Normalized count of sEPSCs | WT | Basal | 1.00 ± 0.00 | 6 | 0.98893 | |
| | | + DCG-IV | 1.00 ± 0.16 | 6 | | |
| | | Basal | 1.00 ± 0.00 | 6 | **0.00039** | |
| | | + WIN | 0.49 ± 0.06 | 6 | | |
| **Figure 6F** **Interneurons** Normalized count of sEPSCs | KO | Basal | 1.00 ± 0.00 | 7 | 0.65232 | WT vs KO, p = 0.75047 |
| | | + DCG-IV | 1.08 ± 0.16 | 7 | | |
| | | Basal | 1.00 ± 0.00 | 7 | **0.00013** | WT vs KO, p = 0.20346 |
| | | +WIN | 0.60 ± 0.05 | 7 | | |
| **Figure 6I** **Mossy cells** Normalized count of sEPSCs | WT | Basal | 1.00 ± 0.00 | 4 | **0.00540** | |
| | | + DCG-IV | 0.48 ± 0.07 | 4 | | |
| | | Basal | 1.00 ± 0.00 | 4 | 0.11715 | |
| | | + WIN | 1.15 ± 0.07 | 4 | | |

WT and KO values were normalized to their respective basal values.

**Figure 7.** Circuit-wide inhibition of Kv7 channels boosted inhibitory drive onto granule cells and corrected granule cell excitability in *Fmr1* KO mice

| Figure 7 | Condition | Genotype | mean ± SEM | Number of cells tested | p value (WT vs KO) | p value (No XE vs + XE991 within genotype) | |
|---|---|---|---|---|---|---|---|
| **Figure 7B** sEPSC frequency (Hz) | Basal | WT | 0.32 ± 0.09 | 7 | 0.76058 | see next two rows | |
| | | KO | 0.29 ± 0.06 | 7 | | | |
| | + XE991 | WT | 0.34 ± 0.05 | 7 | 0.66718 | WT (No XE vs +XE) | 0.71866 |
| | | KO | 0.3 ± 0.07 | 7 | | KO (No XE vs +XE) | 0.75626 |
| **Figure 7C** sIPSC frequency (Hz) | Basal | WT | 1.04 ± 0.12 | 7 | 0.84378 | see next two rows | |
| | | KO | 1.00 ± 0.18 | 7 | | | |
| | + XE991 | WT | 1.08 ± 0.19 | 7 | **0.04061** | WT (No XE vs +XE) | 0.72324 |
| | | KO | 1.96 ± 0.34 | 7 | | KO (No XE vs +XE) | **0.00301** |
| **Figure 7D** E/I ratio (by frequency) | Basal | WT | 0.31 ± 0.07 | 7 | 0.90700 | see next two rows | |
| | | KO | 0.32 ± 0.06 | 7 | | | |
| | + XE991 | WT | 0.37 ± 0.06 | 7 | **0.00590** | WT (No XE vs +XE) | 0.53350 |
| | | KO | 0.15 ± 0.02 | 7 | | KO (No XE vs +XE) | **0.03403** |

| Figure 7 | Step current (pA) | Genotype | mean ± SEM | Number of cells tested | p value (WT vs KO) | p value (No XE vs + XE991 within genotype) |
|---|---|---|---|---|---|---|
| **Figure 7F (Upper panel)** Isolated GCs AP count (Basal) | 50 | WT | 0.60 ± 0.40 | 5 | **0.01695** | see "+XE991" rows |
| | | KO | 3.37 ± 0.83 | 5 | | |
| | 55 | WT | 1.23 ± 0.45 | 5 | **0.00241** | see "+XE991" rows |
| | | KO | 5.84 ± 0.96 | 5 | | |
| | 60 | WT | 2.15 ± 0.65 | 5 | **0.00289** | see "+XE991" rows |
| | | KO | 7.59 ± 1.11 | 5 | | |
| | 65 | WT | 3.56 ± 0.86 | 5 | **0.00332** | see "+XE991" rows |
| | | KO | 9.08 ± 1.03 | 5 | | |
| | 70 | WT | 4.43 ± 0.75 | 5 | **0.00265** | see "+XE991" rows |
| | | KO | 10.60 ± 1.23 | 5 | | |
| | 75 | WT | 5.75 ± 0.85 | 5 | **0.00233** | see "+XE991" rows |
| | | KO | 12.50 ± 1.28 | 5 | | |
| **Figure 7F (Lower panel)** Isolated GCs AP count (+XE991) | 50 | WT | 3.91 ± 1.68 | 5 | **0.04551** | WT (No XE vs +XE) 0.08180 |
| | | KO | 10.21 ± 2.06 | 5 | | KO (No XE vs +XE) **0.03340** |
| | 55 | WT | 6.06 ± 2.05 | 5 | **0.02198** | WT (No XE vs +XE) **0.04124** |
| | | KO | 13.80 ± 1.80 | 5 | | KO (No XE vs +XE) **0.02333** |
| | 60 | WT | 7.47 ± 2.05 | 5 | **0.02318** | WT (No XE vs +XE) **0.02033** |
| | | KO | 15.48 ± 1.99 | 5 | | KO (No XE vs +XE) **0.03574** |
| | 65 | WT | 8.83 ± 2.29 | 5 | **0.01723** | WT (No XE vs +XE) **0.02773** |
| | | KO | 17.66 ± 1.86 | 5 | | KO (No XE vs +XE) **0.02322** |
| | 70 | WT | 9.93 ± 2.28 | 5 | **0.01204** | WT (No XE vs +XE) **0.02826** |
| | | KO | 19.40 ± 1.84 | 5 | | KO (No XE vs +XE) **0.02667** |
| | 75 | WT | 11.38 ± 2.3 | 5 | **0.01001** | WT (No XE vs +XE) **0.01855** |
| | | KO | 21.09 ± 1.76 | 5 | | KO (No XE vs +XE) **0.03012** |
| **Figure 7H (Upper panel)** GCs with intact circuit AP count (Basal) | 50 | WT | 0.79 ± 0.48 | 6 | 0.05186 | see "+XE991" rows |
| | | KO | 3.64 ± 1.20 | 6 | | |
| | 55 | WT | 1.14 ± 0.58 | 6 | **0.02136** | see "+XE991" rows |
| | | KO | 4.94 ± 1.27 | 6 | | |
| | 60 | WT | 2.14 ± 0.78 | 6 | **0.02311** | see "+XE991" rows |
| | | KO | 6.20 ± 1.30 | 6 | | |
| | 65 | WT | 3.04 ± 0.84 | 6 | **0.03448** | see "+XE991" rows |
| | | KO | 7.20 ± 1.47 | 6 | | |
| | 70 | WT | 4.28 ± 0.79 | 6 | **0.03926** | see "+XE991" rows |
| | | KO | 8.35 ± 1.52 | 6 | | |
| | 75 | WT | 5.23 ± 0.73 | 6 | 0.05586 | see "+XE991" rows |
| | | KO | 9.19 ± 1.69 | 6 | | |
| **Figure 7H (Lower panel)** GCs with intact circuit AP count (+XE991) | 50 | WT | 3.90 ± 1.44 | 6 | 0.87305 | WT (No XE vs +XE) **0.03399** |
| | | KO | 4.15 ± 0.54 | 6 | | KO (No XE vs +XE) 0.91875 |
| | 55 | WT | 5.76 ± 1.91 | 6 | 0.93769 | WT (No XE vs +XE) **0.00245** |
| | | KO | 5.60 ± 0.82 | 6 | | KO (No XE vs +XE) 0.35997 |
| | 60 | WT | 6.84 ± 2.05 | 6 | 0.92921 | WT (No XE vs +XE) **0.00665** |
| | | KO | 7.06 ± 1.20 | 6 | | KO (No XE vs +XE) 0.33483 |
| | 65 | WT | 8.27 ± 2.14 | 6 | 0.98096 | WT (No XE vs +XE) **0.00756** |
| | | KO | 8.21 ± 1.29 | 6 | | KO (No XE vs +XE) 0.24645 |
| | 70 | WT | 9.85 ± 2.19 | 6 | 0.85514 | WT (No XE vs +XE) **0.00747** |
| | | KO | 9.36 ± 1.45 | 6 | | KO (No XE vs +XE) 0.20944 |
| | 75 | WT | 10.80 ± 2.22 | 6 | 0.90817 | WT (No XE vs +XE) **0.01147** |
| | | KO | 10.46 ± 1.81 | 6 | | KO (No XE vs +XE) 0.14555 |

**Figure 8.** Circuit-wide inhibition of Kv7 channels restored dentate output during theta-gamma coupling stimulation in *Fmr1* KO mice

| Figure 8 | Condition | Genotype | mean ± SEM | Number of cells tested | p value (WT vs KO) | p value (No XE vs + XE991 within genotype) |
|---|---|---|---|---|---|---|
| **Figure 8C** Excitatory component (norm.) | Basal | WT | 0.45 ± 0.07 | 6 | 0.18795 | see next two rows |
| | | KO | 0.58 ± 0.06 | 7 | | |
| | + XE991 | WT | 0.59 ± 0.06 | 7 | 0.40387 | WT (No XE vs +XE) 0.16120 |
| | | KO | 0.53 ± 0.04 | 8 | | KO (No XE vs +XE) 0.48863 |

| | | | | | | | |
|---|---|---|---|---|---|---|---|
| **Figure 8D** Inhibitory component (norm.) | Basal | WT | 0.12 ± 0.03 | 6 | **0.04190** | see next two rows | |
| | | KO | 0.04 ± 0.02 | 7 | | | |
| | + XE991 | WT | 0.11 ± 0.04 | 7 | 0.98720 | WT (No XE vs +XE) | 0.83913 |
| | | KO | 0.11 ± 0.03 | 8 | | KO (No XE vs +XE) | **0.04970** |
| **Figure 8E** Underlying IPSC (norm.) | Basal | WT | 0.80 ± 0.03 | 6 | **0.00657** | see next two rows | |
| | | KO | 0.62 ± 0.05 | 7 | | | |
| | + XE991 | WT | 0.69 ± 0.04 | 7 | 0.09541 | WT (No XE vs +XE) | 0.06361 |
| | | KO | 0.81 ± 0.05 | 8 | | KO (No XE vs +XE) | **0.01230** |
| **Figure 8F** E/I ratio by peak | Basal | WT | 1.27 ± 0.05 | 6 | **0.01160** | see next two rows | |
| | | KO | 1.69 ± 0.12 | 7 | | | |
| | + XE991 | WT | 1.64 ± 0.16 | 7 | 0.05947 | WT (No XE vs +XE) | 0.05773 |
| | | KO | 1.29 ± 0.09 | 8 | | KO (No XE vs +XE) | **0.03462** |
| **Figure 8G** E/I ratio by charge | Basal | WT | 0.88 ± 0.04 | 6 | **0.00294** | see next two rows | |
| | | KO | 1.35 ± 0.11 | 7 | | | |
| | + XE991 | WT | 1.05 ± 0.09 | 7 | 0.74566 | WT (No XE vs +XE) | 0.11358 |
| | | KO | 1.08 ± 0.05 | 8 | | KO (No XE vs +XE) | **0.03527** |
| **Figure 8H** Excitation window (ms) | Basal | WT | 8.28 ± 0.69 | 6 | **0.00730** | see next two rows | |
| | | KO | 14.03 ± 1.50 | 7 | | | |
| | + XE991 | WT | 9.59 ± 0.67 | 7 | 0.64246 | WT (No XE vs +XE) | 0.20230 |
| | | KO | 9.05 ± 0.89 | 8 | | KO (No XE vs +XE) | **0.04411** |
| **Figure 8K (Upper panel)** granule cell AP probability (Basal) | Control | WT | 0.47 ± 0.03 | 14 | 0.97893 | see "+XE991" rows | |
| | | KO | 0.47 ± 0.03 | 12 | | | |
| | 1st second | WT | 0.09 ± 0.02 | 14 | **0.00060** | see "+XE991" rows | |
| | | KO | 0.21 ± 0.03 | 12 | | | |
| | 2nd second | WT | 0.28 ± 0.03 | 14 | **0.00090** | see "+XE991" rows | |
| | | KO | 0.46 ± 0.04 | 12 | | | |
| | 3rd second | WT | 0.36 ± 0.04 | 14 | **0.01863** | see "+XE991" rows | |
| | | KO | 0.51 ± 0.04 | 12 | | | |
| **Figure 8K (Lower panel)** granule cell AP probability (+XE991) | Control | WT | 0.46 ± 0.02 | 14 | 0.09764 | WT (basal vs XE) | 0.82742 |
| | | KO | 0.39 ± 0.04 | 10 | | KO (basal vs XE) | 0.11948 |
| | 1st second | WT | 0.06 ± 0.01 | 14 | 0.11847 | WT (basal vs XE) | 0.20070 |
| | | KO | 0.10 ± 0.03 | 10 | | KO (basal vs XE) | **0.01054** |
| | 2nd second | WT | 0.23 ± 0.02 | 14 | 0.25639 | WT (basal vs XE) | 0.17289 |
| | | KO | 0.30 ± 0.07 | 10 | | KO (basal vs XE) | **0.04731** |
| | 3rd second | WT | 0.34 ± 0.03 | 14 | 0.76091 | WT (basal vs XE) | 0.85391 |
| | | KO | 0.32 ± 0.06 | 10 | | KO (basal vs XE) | 0.10776 |

**Figure 3—figure supplement 1.** No changes in KCNQ2 and KCNQ3 expression in *Fmr1* KO mice

| Figure 3—figure supplement 1 | Parameter (Western blot) | Genotype | mean ± SEM | Number of mice tested | p value (WT vs KO) |
|---|---|---|---|---|---|
| **Figure 3—figure supplement 1A** | KCNQ2 (whole brain) | WT | 0.69 ± 0.10 | 5 | 0.75314 |
| | | KO | 0.64 ± 0.13 | 5 | |
| **Figure 3—figure supplement 1A** | KCNQ3 (whole brain) | WT | 0.18 ± 0.03 | 5 | 0.15476 |
| | | KO | 0.26 ± 0.03 | 5 | |
| **Figure 3—figure supplement 1C** | KCNQ2 (Dentate gyrus) | WT | 0.29 ± 0.03 | 5 | 0.75101 |
| | | KO | 0.28 ± 0.03 | 5 | |
| **Figure 3—figure supplement 1D** | KCNQ3 (Dentate gyrus) | WT | 0.41 ± 0.14 | 4 | 0.38937 |
| | | KO | 0.49 ± 0.10 | 5 | |

**Figure 4—figure supplement 1.** Changes in miniature synaptic inputs onto mossy cells in *Fmr1* KO mice

| Figure 4—figure supplement 1 | Parameter | Genotype | mean ± SEM | Number of cells tested | p value (WT vs KO) |
|---|---|---|---|---|---|
| **Figure 4—figure supplement 1B** | Cumulative probability* | WT | N/A | 6 | **0.00460** |
| | | KO | N/A | 8 | |
| | mEPSC count (/min) | WT | 709.67 ± 228.51 | 6 | **0.03463** |
| | | KO | 1343.00 ± 154.78 | 8 | |

| | | | | | |
|---|---|---|---|---|---|
| **Figure 4—figure supplement 1C** | mEPSC amplitude (pA) | WT | 72.23 ± 10.86 | 7 | 0.80984 |
| | | KO | 75.46 ± 7.98 | 9 | |
| **Figure 4—figure supplement 1E** | Cumulative probability* | WT | N/A | 18 | 0.10110 |
| | | KO | N/A | 22 | |
| | mIPSC count (/min) | WT | 275.89 ± 33.98 | 18 | 0.30377 |
| | | KO | 325.91 ± 33.31 | 22 | |
| **Figure 4—figure supplement 1F** | mIPSC amplitude (pA) | WT | 32.56 ± 2.83 | 18 | 0.24776 |
| | | KO | 37.49 ± 3.01 | 22 | |

*, K-S test was used for comparison of cumulative probability. N/A, not applicable.

**Figure 5—figure supplement 1.** Changes in miniature synaptic inputs onto hilar interneurons in *Fmr1* KO mice

**Figure 5—figure supplement 1A Number and ratio of interneuron types**

| Genotype \ Celltype | Non-adapting cells | Adapting cells | Burst firing cells |
|---|---|---|---|
| **WT** | 21 (54.1%) | 4 (10.8%) | 12 (35.1%) |
| **KO** | 17 (62.1%) | 3 (10.3%) | 9 (27.6%) |
| **Total (cell type)** | 38 (57.6%) | 7 (10.6%) | 21 (31.8%) |
| **Total (genotype)** | WT 37, KO 29, both genotypes 66 | | |

Chi-square test, $\chi^2$ (2, 66) = 0.02312398 (less than the critical value of 5.9915 for $\alpha$ = 0.05, indicating no significant difference in the cell-type ratios between WT and KO.)

Chi-squre test, **p** = 0.9885046.

| Figure 5—figure supplement 1 | Parameter | Genotype | mean ± SEM | Number of cells tested | p value (WT vs KO) |
|---|---|---|---|---|---|
| **Figure 5—figure supplement 1B** | RMP (mV) | WT | -55.05 ± 0.54 | 37 | 0.52945 |
| | | KO | -54.45 ± 0.83 | 29 | |
| **Figure 5—figure supplement 1C** | Capacitance (pF) | WT | 27.27 ± 1.15 | 37 | 0.68783 |
| | | KO | 28.15 ± 1.98 | 29 | |
| **Figure 5—figure supplement 1D** | Input resistance (MΩ) | WT | 425.49 ± 24.05 | 37 | 0.79275 |
| | | KO | 435.27 ± 28.49 | 29 | |
| **Figure 5—figure supplement 1E** | Threshold (mV) | WT | -39.32 ± 0.52 | 37 | 0.40858 |
| | | KO | -38.68 ± 0.56 | 29 | |
| **Figure 5—figure supplement 1G** | Cumulative probability* | WT | N/A | 8 | **<0.00001** |
| | | KO | N/A | 7 | |
| | mEPSC count (/min) | WT | 534.25 ± 145.85 | 8 | **0.00974** |
| | | KO | 57.86 ± 16.57 | 7 | |
| **Figure 5—figure supplement 1H** | mEPSC amplitude (pA) | WT | 51.69 ± 4.41 | 8 | 0.06326 |
| | | KO | 38.12 ± 5.06 | 7 | |
| **Figure 5—figure supplement 1J** | Cumulative probability* | WT | N/A | 10 | 0.09770 |
| | | KO | N/A | 6 | |
| | mIPSC count (/min) | WT | 61.10 ± 22.02 | 10 | 0.46628 |
| | | KO | 37.33 ± 17.54 | 6 | |
| **Figure 5—figure supplement 1K** | mIPSC amplitude (pA) | WT | 36.33 ± 3.96 | 10 | 0.92243 |
| | | KO | 35.63 ± 6.28 | 6 | |

*, K-S test was used for comparison of cumulative probability. N/A, not applicable.

**Figure 7—figure supplement 1.** Effect of XE991 on spontaneous synaptic inputs onto GCs

| Figure 7—figure supplement 1 | Condition | Genotype | mean ± SEM | Number of cells tested | p value (WT vs KO) | p value (No XE vs + XE991 within genotype) |
|---|---|---|---|---|---|---|
| **Figure 7—figure supplement 1B** sEPSC amplitude (pA) | Basal | WT | 24.10 ± 2.28 | 7 | 0.48259 | see next two rows |
| | | KO | 21.95 ± 1.90 | 7 | | |
| | + XE991 | WT | 19.71 ± 1.55 | 7 | 0.81642 | WT **0.00309** |
| | | KO | 19.07 ± 2.20 | 7 | | KO **0.04308** |
| **Figure 7—figure supplement 1C** sIPSC amplitude (pA) | Basal | WT | 20.27 ± 1.16 | 7 | 0.63095 | see next two rows |
| | | KO | 21.37 ± 1.90 | 7 | | |
| | + XE991 | WT | 17.77 ± 0.95 | 7 | 0.80127 | WT **0.01661** |
| | | KO | 17.30 ± 1.58 | 7 | | KO **0.00189** |

| | | | | | | | |
|---|---|---|---|---|---|---|---|
| **Figure 7—figure supplement 1D** E/I ratio (by amplitude) | Basal | WT | 1.19 ± 0.08 | 7 | 0.08455 | see next two rows | |
| | | KO | 1.03 ± 0.03 | 7 | | | |
| | + XE991 | WT | 1.10 ± 0.04 | 7 | 0.94352 | WT | 0.10015 |
| | | KO | 1.11 ± 0.08 | 7 | | KO | 0.35012 |
| **Figure 7—figure supplement 1E** sEPSC charge transfer (pA·ms) | Basal | WT | 82.92 ± 6.45 | 7 | 0.18959 | see next two rows | |
| | | KO | 69.11 ± 7.56 | 7 | | | |
| | + XE991 | WT | 83.23 ± 5.73 | 7 | 0.37118 | WT | 0.93365 |
| | | KO | 70.78 ± 12.11 | 7 | | KO | 0.82689 |
| **Figure 7—figure supplement 1F** sIPSC charge transfer (pA·ms) | Basal | WT | 73.39 ± 3.98 | 7 | 0.51672 | see next two rows | |
| | | KO | 79.34 ± 7.96 | 7 | | | |
| | + XE991 | WT | 82.31 ± 4.63 | 7 | 0.30529 | WT | 0.08746 |
| | | KO | 72.66 ± 7.74 | 7 | | KO | 0.30306 |
| **Figure 7—figure supplement 1G** E/I ratio (by charge transfer) | Basal | WT | 1.13 ± 0.05 | 7 | 0.15131 | see next two rows | |
| | | KO | 0.98 ± 0.08 | 7 | | | |
| | + XE991 | WT | 1.01 ± 0.03 | 7 | 0.57451 | WT | 0.06407 |
| | | KO | 0.95 ± 0.09 | 7 | | KO | 0.88906 |

**Figure 7—figure supplement 2.** Estimation of direct and circuit effects of XE991 on granule cell excitability

| **Figure 7—figure supplement 2** | **Step current (pA)** | **Genotype** | **Values estimated from mean** |
|---|---|---|---|
| **Figure 7—figure supplement 2A** XE991 action within granule cells (number of APs) | 50 | WT | 3.31 |
| | | KO | 6.84 |
| | 55 | WT | 4.83 |
| | | KO | 7.96 |
| | 60 | WT | 5.32 |
| | | KO | 7.89 |
| | 65 | WT | 5.27 |
| | | KO | 8.58 |
| | 70 | WT | 5.50 |
| | | KO | 8.80 |
| | 75 | WT | 5.63 |
| | | KO | 8.59 |
| **Figure 7—figure supplement 2B** XE991 action from dentate circuit (number of APs) | 50 | WT | 0.03 |
| | | KO | -6.68 |
| | 55 | WT | 0.38 |
| | | KO | -6.25 |
| | 60 | WT | 0.81 |
| | | KO | -5.96 |
| | 65 | WT | 1.17 |
| | | KO | -5.99 |
| | 70 | WT | -0.10 |
| | | KO | -5.76 |
| | 75 | WT | 0.13 |
| | | KO | -4.90 |

**Figure 8—figure supplement 2.** Circuit-wide inhibition of Kv7 channels enhanced the gamma burst-induced suppression of EPSP integration in *Fmr1* KO mice

| **Figure 8—figure supplement 2** | **Parameter** | **Genotype** | **mean ± SEM** | **Number of cells tested** | **p value (WT vs KO)** | **p value (Basal vs +XE991 within genotype)** | |
|---|---|---|---|---|---|---|---|
| **Figure 8—figure supplement 2B** | EPSP (mV) (No XE991) | WT | 22.65 ± 1.10 | 14 | 0.16073 | see next tow rows | |
| | | KO | 20.66 ± 0.73 | 12 | | | |
| **Figure 8—figure supplement 2D** | EPSP (mV) (+ XE991) | WT | 20.89 ± 0.57 | 14 | 0.32137 | WT (No XE vs +XE) | 0.16896 |
| | | KO | 20.04 ± 0.57 | 10 | | KO (No XE vs +XE) | 0.52682 |

| Figure 8—figure supplement 2 | Stimulus # | Genotype | mean ± SEM | Number of cells tested | p value (WT vs KO) |
|---|---|---|---|---|---|
| **Figure 8—figure supplement 2B (Left panel)** EPSP normalized amplitude (control stimulus) (No XE991) | 1 | WT | 1.04 ± 0.01 | 14 | 0.59725 |
| | | KO | 1.03 ± 0.01 | 12 | |
| | 2 | WT | 1.01 ± 0.02 | 14 | 0.51456 |
| | | KO | 1.03 ± 0.02 | 12 | |
| | 3 | WT | 1.01 ± 0.02 | 14 | 0.98511 |
| | | KO | 1.01 ± 0.01 | 12 | |
| | 4 | WT | 1.03 ± 0.02 | 14 | 0.46543 |
| | | KO | 1.04 ± 0.02 | 12 | |
| | 5 | WT | 1.01 ± 0.01 | 14 | 0.71914 |
| | | KO | 1.01 ± 0.02 | 12 | |
| | 6 | WT | 1.00 ± 0.01 | 14 | 0.21751 |
| | | KO | 1.04 ± 0.03 | 12 | |
| | 7 | WT | 1.00 ± 0.01 | 14 | 0.45138 |
| | | KO | 1.01 ± 0.02 | 12 | |
| | 8 | WT | 1.00 ± 0.02 | 14 | 0.49648 |
| | | KO | 0.98 ± 0.02 | 12 | |
| | 9 | WT | 1.00 ± 0.02 | 14 | 0.13733 |
| | | KO | 0.96 ± 0.02 | 12 | |
| | 10 | WT | 1.00 ± 0.01 | 14 | 0.52223 |
| | | KO | 0.99 ± 0.02 | 12 | |
| | 11 | WT | 0.99 ± 0.01 | 14 | 0.97942 |
| | | KO | 0.99 ± 0.02 | 12 | |
| | 12 | WT | 0.98 ± 0.01 | 14 | 0.80630 |
| | | KO | 0.97 ± 0.01 | 12 | |
| | 13 | WT | 1.00 ± 0.01 | 14 | 0.94782 |
| | | KO | 1.00 ± 0.02 | 12 | |
| | 14 | WT | 0.98 ± 0.01 | 14 | 0.68896 |
| | | KO | 0.97 ± 0.01 | 12 | |
| | 15 | WT | 1.00 ± 0.01 | 14 | 0.29475 |
| | | KO | 0.99 ± 0.01 | 12 | |
| **Figure 8—figure supplement 2B (Right panel)** EPSP normalized amplitude (Test stimulus) (NO XE991) | 1 | WT | 0.46 ± 0.03 | 14 | **<0.00001** |
| | | KO | 0.74 ± 0.04 | 12 | |
| | 2 | WT | 0.64 ± 0.02 | 14 | **<0.00001** |
| | | KO | 0.89 ± 0.04 | 12 | |
| | 3 | WT | 0.74 ± 0.02 | 14 | **<0.00001** |
| | | KO | 0.98 ± 0.04 | 12 | |
| | 4 | WT | 0.78 ± 0.02 | 14 | **0.00002** |
| | | KO | 1.01 ± 0.04 | 12 | |
| | 5 | WT | 0.76 ± 0.02 | 14 | **0.00003** |
| | | KO | 1.01 ± 0.05 | 12 | |
| | 6 | WT | 0.80 ± 0.02 | 14 | **0.00003** |
| | | KO | 1.02 ± 0.04 | 12 | |
| | 7 | WT | 0.80 ± 0.02 | 14 | **0.00005** |
| | | KO | 1.01 ± 0.04 | 12 | |
| | 8 | WT | 0.79 ± 0.03 | 14 | **0.00013** |
| | | KO | 1.01 ± 0.04 | 12 | |
| | 9 | WT | 0.81 ± 0.02 | 14 | **<0.00001** |
| | | KO | 1.03 ± 0.04 | 12 | |
| | 10 | WT | 0.80 ± 0.02 | 14 | **0.00005** |
| | | KO | 1.03 ± 0.04 | 12 | |
| | 11 | WT | 0.81 ± 0.02 | 14 | **0.00010** |
| | | KO | 1.02 ± 0.04 | 12 | |
| | 12 | WT | 0.83 ± 0.02 | 14 | **0.00002** |
| | | KO | 1.05 ± 0.04 | 12 | |
| | 13 | WT | 0.80 ± 0.02 | 14 | **0.00115** |
| | | KO | 0.98 ± 0.05 | 12 | |
| | 14 | WT | 0.80 ± 0.03 | 14 | **0.00019** |
| | | KO | 1.02 ± 0.04 | 12 | |

| | | | | | |
|---|---|---|---|---|---|
| | 15 | WT | 0.81 ± 0.03 | 14 | **0.00010** |
| | | KO | 1.02 ± 0.04 | 12 | |
| **Figure 8—figure supplement 2D (Left panel)** EPSP normalized amplitude (control stimulus) (+ XE991) | 1 | WT | 1.03 ± 0.01 | 14 | 0.07020 |
| | | KO | 0.99 ± 0.01 | 10 | |
| | 2 | WT | 1.00 ± 0.02 | 14 | 0.55476 |
| | | KO | 0.98 ± 0.02 | 10 | |
| | 3 | WT | 1.01 ± 0.02 | 14 | 0.98112 |
| | | KO | 1.01 ± 0.01 | 10 | |
| | 4 | WT | 1.02 ± 0.01 | 14 | 0.71698 |
| | | KO | 1.01 ± 0.02 | 10 | |
| | 5 | WT | 1.01 ± 0.02 | 14 | 0.13496 |
| | | KO | 0.98 ± 0.01 | 10 | |
| | 6 | WT | 0.97 ± 0.02 | 14 | 0.17428 |
| | | KO | 1.00 ± 0.02 | 10 | |
| | 7 | WT | 1.00 ± 0.01 | 14 | 0.09702 |
| | | KO | 1.02 ± 0.01 | 10 | |
| | 8 | WT | 1.00 ± 0.01 | 14 | 0.63306 |
| | | KO | 1.01 ± 0.02 | 10 | |
| | 9 | WT | 1.01 ± 0.02 | 14 | 0.67985 |
| | | KO | 1.02 ± 0.01 | 10 | |
| | 10 | WT | 1.00 ± 0.01 | 14 | 0.80769 |
| | | KO | 1.00 ± 0.02 | 10 | |
| | 11 | WT | 1.00 ± 0.01 | 14 | 0.93354 |
| | | KO | 1.00 ± 0.02 | 10 | |
| | 12 | WT | 1.00 ± 0.01 | 14 | 0.64994 |
| | | KO | 0.99 ± 0.03 | 10 | |
| | 13 | WT | 0.97 ± 0.02 | 14 | 0.31199 |
| | | KO | 0.99 ± 0.02 | 10 | |
| | 14 | WT | 0.99 ± 0.01 | 14 | 0.96013 |
| | | KO | 0.99 ± 0.02 | 10 | |
| | 15 | WT | 0.99 ± 0.01 | 14 | 0.15667 |
| | | KO | 1.01 ± 0.01 | 10 | |
| **Figure 8—figure supplement 2D (Right panel)** EPSP normalized amplitude (Test stimulus) (+ XE991) | 1 | WT | 0.71 ± 0.03 | 14 | 0.08684 |
| | | KO | 0.62 ± 0.04 | 10 | |
| | 2 | WT | 0.77 ± 0.03 | 14 | **0.02263** |
| | | KO | 0.66 ± 0.03 | 10 | |
| | 3 | WT | 0.82 ± 0.04 | 14 | 0.06573 |
| | | KO | 0.72 ± 0.03 | 10 | |
| | 4 | WT | 0.86 ± 0.04 | 14 | 0.10039 |
| | | KO | 0.76 ± 0.04 | 10 | |
| | 5 | WT | 0.87 ± 0.03 | 14 | 0.15131 |
| | | KO | 0.79 ± 0.04 | 10 | |
| | 6 | WT | 0.87 ± 0.03 | 14 | 0.10975 |
| | | KO | 0.78 ± 0.05 | 10 | |
| | 7 | WT | 0.89 ± 0.04 | 14 | 0.10782 |
| | | KO | 0.80 ± 0.04 | 10 | |
| | 8 | WT | 0.89 ± 0.03 | 14 | 0.05007 |
| | | KO | 0.79 ± 0.03 | 10 | |
| | 9 | WT | 0.86 ± 0.04 | 14 | 0.22119 |
| | | KO | 0.79 ± 0.04 | 10 | |
| | 10 | WT | 0.91 ± 0.04 | 14 | 0.08052 |
| | | KO | 0.80 ± 0.05 | 10 | |
| | 11 | WT | 0.89 ± 0.04 | 14 | 0.29036 |
| | | KO | 0.83 ± 0.04 | 10 | |
| | 12 | WT | 0.90 ± 0.04 | 14 | 0.30202 |
| | | KO | 0.83 ± 0.05 | 10 | |
| | 13 | WT | 0.90 ± 0.04 | 14 | 0.29607 |
| | | KO | 0.84 ± 0.04 | 10 | |
| | 14 | WT | 0.92 ± 0.03 | 14 | 0.12978 |
| | | KO | 0.84 ± 0.03 | 10 | |
| | 15 | WT | 0.91 ± 0.04 | 14 | 0.26870 |
| | | KO | 0.85 ± 0.03 | 10 | |
